# Nucleosome reorganisation in breast cancer tissues

**DOI:** 10.1101/2023.04.17.537031

**Authors:** Divya R. Jacob, Wilfried M. Guiblet, Hulkar Mamayusupova, Mariya Shtumpf, Isabella Ciuta, Luminita Ruje, Svetlana Gretton, Milena Bikova, Clark Correa, Emily Dellow, Shivam P. Agrawal, Navid Shafiei, Anastasija Drobysevskaja, Chris M. Armstrong, Jonathan D. G. Lam, Yevhen Vainshtein, Christopher T. Clarkson, Graeme J. Thorn, Kai Sohn, Madapura M. Pradeepa, Sankaran Chandrasekharan, Greg N. Brooke, Elena Klenova, Victor B. Zhurkin, Vladimir B. Teif

## Abstract

Nucleosome repositioning in cancer is believed to cause many changes in genome organisation and gene expression. Understanding these changes is important to elucidate fundamental aspects of cancer. It is also important for medical diagnostics based on cell-free DNA (cfDNA), which originates from genomic DNA regions protected from digestion by nucleosomes. Here we have generated high resolution nucleosome maps in paired tumour and normal tissues from the same breast cancer patients using MNase-assisted histone H3 ChIP-seq and compared them with the corresponding cfDNA from blood plasma. This analysis has detected single-nucleosome repositioning at key regulatory regions in a patient-specific manner and common cancer-specific patterns across patients. The nucleosomes gained in tumour versus normal tissue were particularly informative of cancer pathways, with ∼20-fold enrichment at CpG islands, a large fraction of which marked promoters of genes encoding DNA-binding proteins. In addition, tumour tissues were characterised by a 5-10 bp decrease in the average distance between nucleosomes (nucleosome repeat length, NRL), which is qualitatively similar to the differences between pluripotent and differentiated cells. These effects were correlated with gene activity, DNA sequence repeats abundance, differential DNA methylation and binding of linker histone variants H1.4 and H1X. Our findings provide a new mechanistic understanding of nucleosome repositioning in tumour tissues that can be valuable for patient stratification and monitoring using liquid biopsies.

## Introduction

Nucleosome positioning is one of the main determinants of gene regulation. Apart from its fundamental importance, understanding the consequences of nucleosome positioning is promising for clinical diagnostics based on cell-free DNA (cfDNA) (Shtumpf et al. 2022). cfDNA is composed of DNA fragments released into body fluids after digestion of nuclear chromatin by apoptotic nucleases present *in situ*. Such DNA fragments are protected from nuclease digestion by nucleosomes and other DNA-bound proteins, and therefore cfDNA analysis allows the construction of nucleosome maps. The natural process of cfDNA formation is similar to MNase-seq experiments in which chromatin is digested between nucleosomes by micrococcal nuclease (MNase), and MNase-resistant DNA fragments are analysed (Teif et al. 2012; Snyder et al. 2016). While the connection between nucleosome positioning in tissues and cfDNA is of particular importance, only a few low- to moderate-resolution nucleosome landscapes have been previously reported in breast cell lines (Lidor Nili et al. 2010; Shimbo et al. 2013; Bansal et al. 2015; Lavender et al. 2016; Takaku et al. 2016; Cole et al. 2021; Yang et al. 2021), and none available in paired normal/tumour primary tissues from breast cancer (BRC) patients. It is worth noting that the lack of nucleosome positioning datasets in paired normal/tumour tissues is also characteristic for other cancers.

Our recent study in chronic lymphocytic leukaemia uncovered some of the rules of cancer-specific nucleosome repositioning in B-cells from peripheral blood (Piroeva et al. 2023), but nucleosome repositioning in solid tissues remains enigmatic. One of the reasons for this is that chromatin in solid tissues may have vastly different properties than chromatin from immortalised cell lines, which have been used historically as model systems for nucleosome positioning. For example, as early as 1976, Compton et al (Compton et al. 1976) compared nucleosome repeat length (NRL) values from different studies in normal and cancer cells and observed that normal cells in culture have shorter NRLs in comparison with primary cells in tissues. This means that the detection of differences in nucleosome positioning between normal and tumour cells in culture may be challenging, and it may not represent the effects happening *in situ*. To overcome these challenges, here we have generated the first resource comprised of ultra-high resolution nucleosome maps in paired normal and tumour breast tissues, as well as in cfDNA from the same patients.

The dataset reported here contains about 20 billion paired-end sequenced reads, which makes it one of the largest nucleosome positioning resources of its kind. This allowed us to perform a systematic analysis of BRC-specific nucleosome repositioning *in situ*. Discrete nucleosome repositioning was enriched at gene promoters encoding key BRC transcription factors (TFs), as well as at cancer-sensitive subsets of TF binding sites. In addition, we found that cancer tissues are characterised by a 5-10 bp decrease in NRL and we have investigated potential molecular mechanisms of such dramatic shortening of distances between nucleosomes. These new results open clinically relevant avenues based on analyses of cancer-specific nucleosome reorganisation.

## Results

### Cancer-specific nucleosome repositioning in breast tissues

We determined ultra-high resolution nucleosome positioning landscapes in paired normal and tumour breast tissues from four BRC patients using MNase-seq and MNase-assisted histone H3 ChIP-seq (MNase- H3) (Figure 1A). The high sequencing coverage allowed us to define changes at the level of single nucleosomes (Figure 1B). First, for each of the two conditions (normal and tumour), we defined stable nucleosomes as regions protected from MNase digestion in a given condition in a given patient and confirmed in the same condition in at least one other patient from the cohort studied here. Then lost/gained nucleosomes were defined as those which are stable in one condition as defined above and do not overlap with any stable nucleosomes in the other condition. As a result of this definition, we were able to arrive at a set of ∼50,000 gained/lost nucleosomes using Patient P2 as an example (see Materials and Methods).

**Figure 1.**
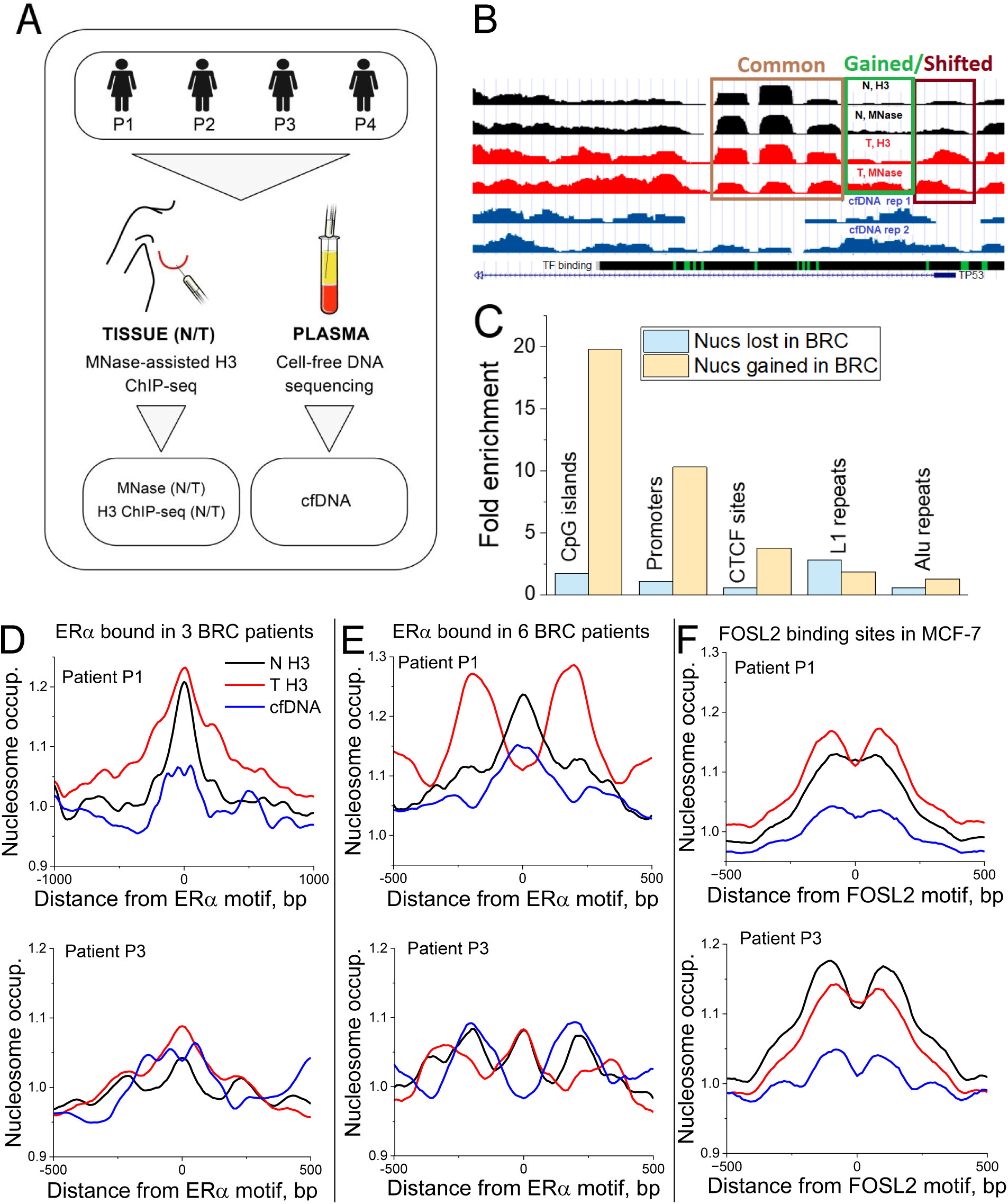
The study design and small-scale nucleosome repositioning analysis. A) Scheme of the study: paired tumour/normal breast tissues taken from breast cancer patients numbered P1-P4 were used to determine nucleosome positioning using MNase-seq and MNase-assisted histone H3 ChIP-seq (MNase-H3), complemented by whole-genome sequencing of cell-free DNA extracted from blood plasma from the same patients. The analysis of each sample was performed individually, without pooling. B) Example genomic region (Chr 17: 7,855,262-7,592,516) enclosing gene TP53 and nucleosome occupancy maps for patient P3. Tracks top to bottom: normal tissue MNase-H3 and MNase-seq; tumour tissue MNase-H3 and MNase-seq; cfDNA in two replicates; binding sites of 161 TFs derived from ChIP-seq from ENCODE with Factorbook motifs. Rectangles show regions containing common (brown), gained (green) and shifted nucleosomes (violet). The gained and shifted classifications are not mutually exclusive. C) Fold enrichment of nucleosomes that were gained and lost in all tumours versus healthy breast tissues in CpG islands, promoters, CTCF binding sites, L1 repeats and Alu repeats, in comparison with the intersections with these regions expected by chance. In all cases except for nucleosomes lost in BRC at CTCF sites, enrichments are statistically significant (Fisher test, p < 0.005). D, E, F) Representative nucleosome occupancy profiles around TF binding sites. D) and E) Nucleosome profiles in patient P1 (top) and patient P3 from the current cohort (bottom) around ERα sites bound in any three out of six BRC patients from the cohort of Severson et al., 2018 (D) and in sites bound by ERα in all six BRC patients from Severson et al., 2018 (E) (10-bp smoothing). F) Nucleosome profiles in patient P1 (top) and patient P3 (bottom) around FOSL2 binding sites determined in MCF-7 cells. Black lines – tumour tissues; red – normal tissues; blue – cfDNA from the same patient.

The nucleosomes gained in tumour tissues defined above were 19-fold enriched at CpG islands and 10-fold enriched at promoters, including promoters of 316 genes encoding DNA-binding proteins (for both effects *p* < 1e-100, Fisher Test) (Figure 1C). Gained nucleosomes were also prevalent at CTCF binding sites. These changes in the nucleosome landscape may define differential accessibility of DNA to regulatory proteins, underlying dysregulated cancer-specific pathways. Gained nucleosomes were particularly predictive of breast cancer pathways. The network of genes encoding DNA-binding proteins, marked by gained nucleosomes at their promoters, was significantly enriched for protein-protein interactions (*p* < 1e-16), with major nodes represented by known BRC-associated TFs such as RXRA, PPARγ, CREBBP, etc. (Supplementary Figure S2). On the other hand, nucleosomes which were stably present in healthy breast tissues but lost in tumour did not show such enrichment in cis-regulatory regions and were instead modestly enriched inside L1 repeats (Figure 1C).

### Nucleosome changes at sub-classes of transcription factor (TF) binding sites

To complement the above analysis of promoter nucleosome occupancy, we also investigated nucleosome patterns determined in the current patient cohort at specific TF binding sites confirmed by published ChIP-seq datasets in other BRC patients. One of the most important TFs involved in breast cancer and used for its classification/stratification is Estrogen Receptor alpha (ERα)(Brett et al. 2021). We observed cancer-specific nucleosome changes at two classes of ERα binding sites that occurred in a patient-specific fashion, as exemplified in Figure 1D-E. First, a class of less stringent ERα binding sites, defined as a consensus dataset of ERα ChIP-seq peaks present in any three out of six BRC patients from the study of Severson et al (Severson et al. 2018), did not show significant changes in nucleosome occupancy between normal and tumour breast tissues and cfDNA of patients P1 and P3 from our study (Figure 1D). Second, a class of more stringent binding sites occupied by ERα in all six patients from the cohort of Severson et al (Severson et al. 2018) was characterised by pronounced nucleosome changes between normal/tumour breast tissues and cfDNA in patients from our cohort (Figure 1E). Patient P1, who has clinically confirmed ER-positive status, showed a clear change in the shape of the nucleosome profile between tumour and adjacent normal breast tissue (nucleosome depletion at the centres of ERα binding sites and oscillations of nucleosome occupancy around ERα sites in the tumour, as opposed to a peak of nucleosome occupancy at ERα binding sites in normal tissue). On the other hand, patient P3, for whom our immunoblotting did not detect the presence of ERα (Supplementary Figure S1), was characterised by similar shapes of nucleosome occupancy profiles at stringent ERα sites in both paired normal and tumour tissues. Interestingly, the cfDNA profile of patient P1 resembled that of normal tissue, while the cfDNA profile of patient P3 had an inverted shape in comparison with both normal and tumour breast tissues from this patient. Patient-specific changes were also specific to other TFs, although not as pronounced as for ERα, as exemplified by FOSL2 binding sites (Figure 1F). This analysis suggests that not all experimentally confirmed TF binding sites are equally suitable as diagnostic markers, and sub-selection of such sites may play a particularly important role. Thus, a systematic screening of TF binding sites for the potential roles of diagnostics markers deserves a separate study.

### Nucleosome repeat length shortening in breast cancer

Next, we asked whether there is some general change of nucleosome organisation in BRC that affects all patients. We calculated the distribution of genomic distances between centres of neighbouring nucleosomes, which revealed a dramatic shift towards smaller distances in cancer in comparison with adjacent non-cancer tissues from the same patients (Figure 2A). This resulted in a significant decrease in the average genome-wide distance between neighbouring nucleosomes (the Nucleosome Repeat Length, NRL) in tumour versus normal tissues. This effect of genome-wide NRL shortening by 5-10 bp in cancer was observed in all MNase-H3 samples from this study and in MNase-seq for three out of four patients (Figure 2B and Supplementary Table S5). The only outlier was the “Normal” MNase-seq sample of patient P1. However, paired MNase-H3 samples from the same patient P1 were consistent with NRL shortening in other patients. Thus, the effect of NRL shortening was detected in all patients in this cohort. Overall, genome-wide NRL decrease in tumour tissues was statistically significant (*p* = 0.014, paired sample *t*-test). A number of previous works characterised NRL changes in cell differentiation (Compton et al. 1976; Valouev et al. 2011; Teif et al. 2017; Willcockson et al. 2021; Yusufova et al. 2021), and we have shown that highly proliferating embryonic stem cells have shorter NRL than their differentiated counterparts (Teif et al. 2012). However, to the best of our knowledge, this is the first report of a decrease of NRL in tumour versus adjacent non-tumour breast tissue from the same patients. cfDNA from blood plasma of BRC patients was characterised by NRL values similar to those in tumour tissues (Supplementary Table S6).

**Figure 2.**
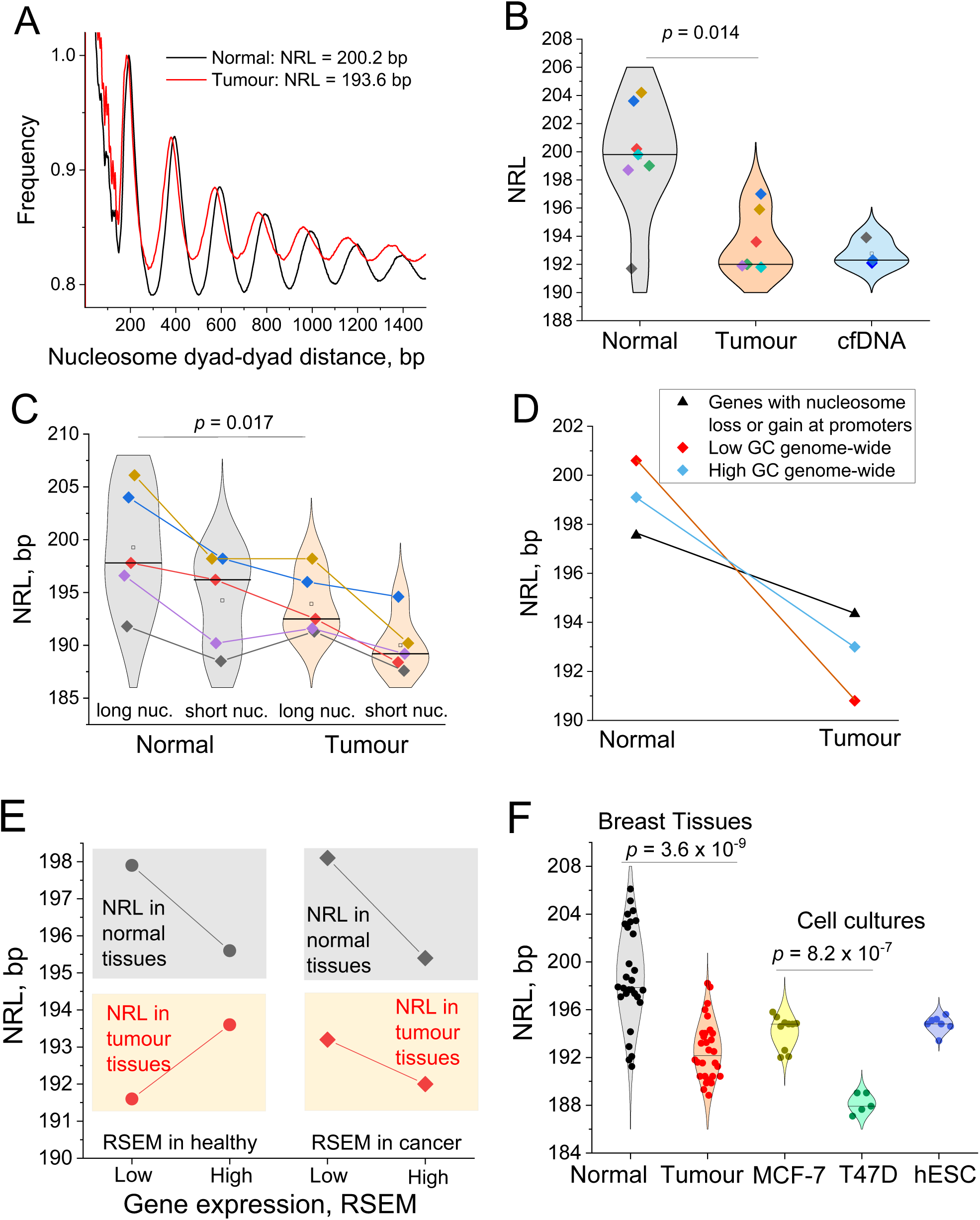
Nucleosome repeat length shortening in cancer. A) Frequencies of distances between dyads of DNA fragments protected from MNase digestion in paired normal and tumour breast tissue samples from patient P2. B) Genome-wide NRL values calculated for each sample reported in this study. The difference of NRLs between normal and tumour tissues is statistically significant (p = 0.014, paired sample t-test). Open squares – average values across all samples in a given condition; horizontal lines – median values for a given condition. The colours of the points corresponding to NRL values in N/T tissues are assigned per patient and per experiment type as follows: MNase-seq: patient P1 (black), P2 (red), P3 (blue), P4 (green); MNase-assisted H3 ChIP-seq: patient P1 (violet), P3 (brown), P4 (cyan). Panels (A) and (B) use DNA fragments of sizes 120-180 bp. C) NRL in normal (grey violins) and tumour (beige violins) breast tissue calculated inside genomic regions enriched (“long nuc.”) and depleted with DNA fragments of sizes 160-180 bp (“short nuc”). The colours of the points are assigned per patient and per experiment type in the same way as in (B). Paired-sample t-test p-values are indicated on the figure. Open squares – average values across all samples in a given condition; horizontal lines – median values for a given condition. D) NRL in normal and tumour breast tissue samples from patient P2 inside genes which are marked by nucleosome loss or gain at their promoters (black), compared with NRL calculated genome-wide in 10 kb regions with low GC content (<40%) (red) and high GC content (>40%) (blue). E) NRL in normal (black) and tumour breast tissues from patient P2 from this study (red), calculated inside genes which are highly- or lowly-expressed in a healthy state (circles) or breast cancer (squares). This analysis is based on 3,000 genes with largest and smallest RSEM values in normal/cancer cells reported by the TCGA Cancer Atlas. F) NRL in MCF-7 and T47D breast cancer cells as well as human embryonic stem cells (hESC), compared with NRL in normal and tumour breast tissues. For T47D and hESC, each point corresponds to one replicate experiment. For MCF-7, each point corresponds to a sample with different MNase digestion level. For breast tissues, each point corresponds to one sample from this work, in a subset of genomic regions enriched with DNA fragment sizes in one of the following ranges [100-120 bp], [120-140 bp], [140-160 bp], [160-180 bp], [180-200 bp].

### The NRL decrease in tumours is modulated by differential gene expression, chromatin digestion levels, and local GC content, in a patient-specific manner

What can potentially explain the large NRL changes that we detected? Firstly, we asked whether the NRL shortening in breast cancer is related to differences in the sizes of DNA fragments protected from MNase digestion. The sizes of DNA fragments protected from MNase digestion (hereafter referred to as “DNA fragments”) may depend on the experimental setup, and therefore it is important to uncouple these effects from NRL changes that reflect the chromatin biology. It’s worth noting that the effect of cfDNA fragment size shortening in cancer is already extensively used in liquid biopsies (Shtumpf et al. 2022), but the effect of NRL shortening has been unknown so far. Typical nucleosome sizes are around 147 bp, while the nucleosome plus DNA linker (termed chromatosome), protects around 160-180 bp. Figure 2C shows that genomic regions enriched with chromatosome-size DNA fragments (160-180 bp, denoted as “long nucleosomes”) have a longer NRL, while those depleted of 160-180 bp fragments have a smaller NRL. However, for both types of regions tumour had a shorter NRL than normal tissue. Thus, cancer-specific NRL shortening cannot be explained by the level of chromatin digestion *per se*.

Next, we asked whether the effect of the NRL decrease is more pronounced at specific genes. The comparison of the NRL in genes marked by nucleosome repositioning at their promoters did not reveal significant differences between those associated with gained versus lost nucleosomes (Figure 2D). Furthermore, the cancer-specific NRL decrease inside genes marked by differential nucleosome occupancy was smaller than the genome-wide change (Figure 2D). The genome-wide NRL decrease in cancer was more pronounced in regions with low GC content ([GC] < 40%, Figure 2D), which is generally outside genes, since most genes have a GC content higher than 45%.

Regarding intragenic NRL changes, we made a comparison between highly- and lowly-expressed genes (Figure 2E). Previous studies suggested that the NRL is decreased over more active genes (Valouev et al. 2011; Voong et al. 2016; Baldi et al. 2018b). When we stratified genes based on their expression in cancer using the TCGA cancer genome atlas (Hoadley et al. 2018), the NRL was indeed smaller for highly- than lowly-expressed genes. When gene expression was stratified based on healthy controls, the NRL measured in healthy tissues was also smaller inside highly-expressed genes, but the NRL measured in tumour tissues did not follow this rule. Altered gene expression may have changed the NRL inside these genes in tumour tissue. Importantly, this effect was in all cases smaller than the major effect of the NRL decrease in cancer tissues versus normal tissues.

The global factors defining the NRL decrease in cancer cells can be compared with those acting in stem cells, which have a smaller NRL than differentiated cells (Teif et al. 2012). Indeed, human embryonic stem cells (hESCs) have an NRL which is more similar to tumour tissue than to normal breast tissues (Figure 2F and Supplementary Table S7). Our calculation showed that breast cancer cell lines MCF-7 and T47D also have smaller NRLs resembling those in tumour tissues (Figure 2F and Supplementary Tables S8-9). Interestingly, T47D had an even smaller NRL in comparison with MCF-7, which was robustly reproducible using eight MCF-7 datasets with different levels of MNase digestion and five replicate experiments in T47D (see Supplementary Materials and Methods). Thus, NRL shortening in breast cancer is a general effect.

### Dependence of nucleosome sensitivity to MNase digestion in tumours on GC content and DNA methylation

To study additional properties of nucleosome repositioning in BRC, we classified nucleosomes into “common nucleosomes” (stable MNase-protected DNA fragments of a given patient found at the same locations in both normal and tumour tissues) and “shifted nucleosomes” (stable MNase-protected DNA fragments of a given patient based on normal tissue that changed their genomic coordinates >20% in tumour). Shifted nucleosomal DNA fragments included a large subnucleosomal fraction (120-140 bp) and were on average shorter than the stable nucleosomes which did not change their locations in cancer (Figure 3A). This may correspond to partial nucleosome unwrapping or nucleosomes with incomplete histone octamer. Common nucleosomes were characterised by DNA methylation enrichment, while shifted nucleosomes were depleted of DNA methylation (Figure 3B). This is consistent with previous reports that DNA methylation stabilises nucleosomes while de- methylation or hydroxymethylation makes nucleosomes more MNase-sensitive (Teif et al. 2014; Wiehle et al. 2019; Teif et al. 2020).

**Figure 3.**
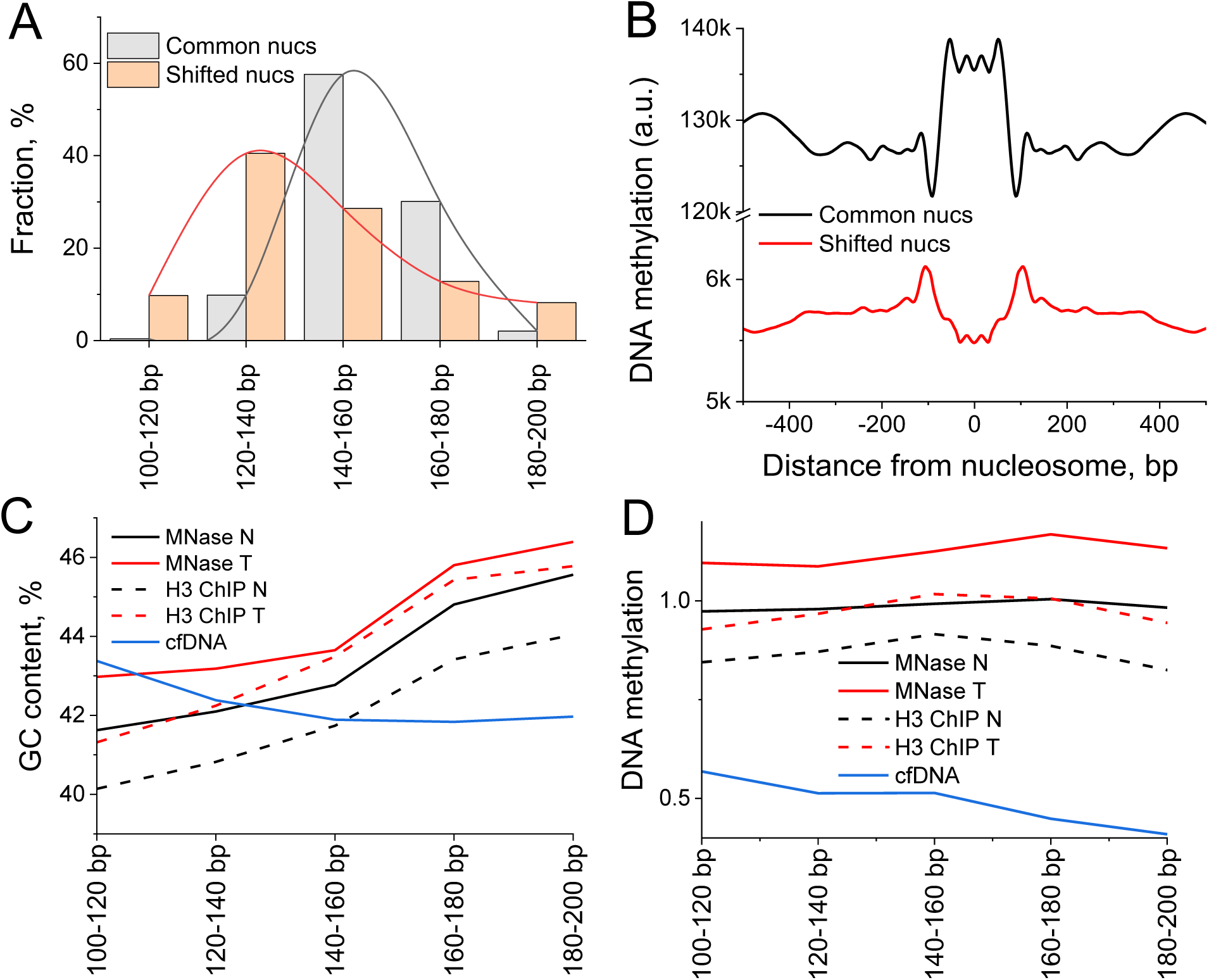
Interplay between nucleosome stability in BRC, DNA fragment sizes, GC content and DNA methylation. A) The distribution of sizes of common nucleosomes that did not shift in cancer (grey) is skewed towards larger sizes in comparison with shifted nucleosomes whose coordinates changed >20% in cancer (red). B) Aggregate DNA methylation profile based on MCF-7 cells plotted around common and shifted nucleosomes determined in this study. C) GC content of DNA fragments protected from digestion in normal (black) and cancer breast tissues (red) and cfDNA from the same patient cohort (blue). D) DNA methylation based on MCF-7 cells averaged across DNA fragments of different sizes protected from MNase digestion from panel (C).

The overall distribution of DNA fragments protected from MNase digestion was more enriched with shorter sizes in cancer in comparison with normal samples (Supplemental Figure S3). This distribution appeared to correlate with the GC content, with longer fragments being more GC-rich (Figure 3C and Supplemental Figures S4D). This is in line with known effects on DNA fragment lengths of the level of MNase digestion (Teif et al. 2014; Chereji et al. 2016). Interestingly, the dependence of the fragment length on DNA methylation was different in breast tissues versus cfDNA (Figure 3D). The size distribution of DNA fragments protected from MNase digestion was similar inside active and inactive chromatin domains (A- and B-domains), repetitive and non-repetitive regions, separated into *Alu*, LINE1, alpha-satellite and micro-satellite repeats (Supplementary Figure S4A-C) and inside regions marked by H3K9me2/H3K9me3 in breast cell lines (Supplementary Figure S5). The GC profile around DNA fragments protected from digestion showed larger differences between normal and cancer samples in the case of shorter fragment sizes, which was the case both for breast tissues and cfDNA (Supplementary Figure S6).

### Relationship between nucleosome reorganisation in BRC and linker histones

Linker histones are among the most abundant chromatin proteins that can affect nucleosome organisation. To investigate how DNA protection by linker histones might differ between healthy and tumour cells, we have used recently reported maps of H1 histone variants determined with ChIP-seq in the BRC cell line T47D (Serna-Pujol et al. 2022). Genome-wide, the occupancy of linker histone variants H1.0, H1.2 and H1.5 is negatively correlated with GC content, while histones H1.4 and H1X are preferentially bound to GC-rich regions (Figure 4A). By comparing H1 maps with the nucleosome maps determined in this study, we clarified that histones H1.0, H1.2 and H1.5 were depleted near nucleosome dyads for protected DNA fragments of all sizes, while H1.4 and H1X occupancies were diminished for sub-nucleosomal and nucleosome-size DNA fragments (<160 bp) but increased for larger-size fragments (Figure 4B).

**Figure 4.**
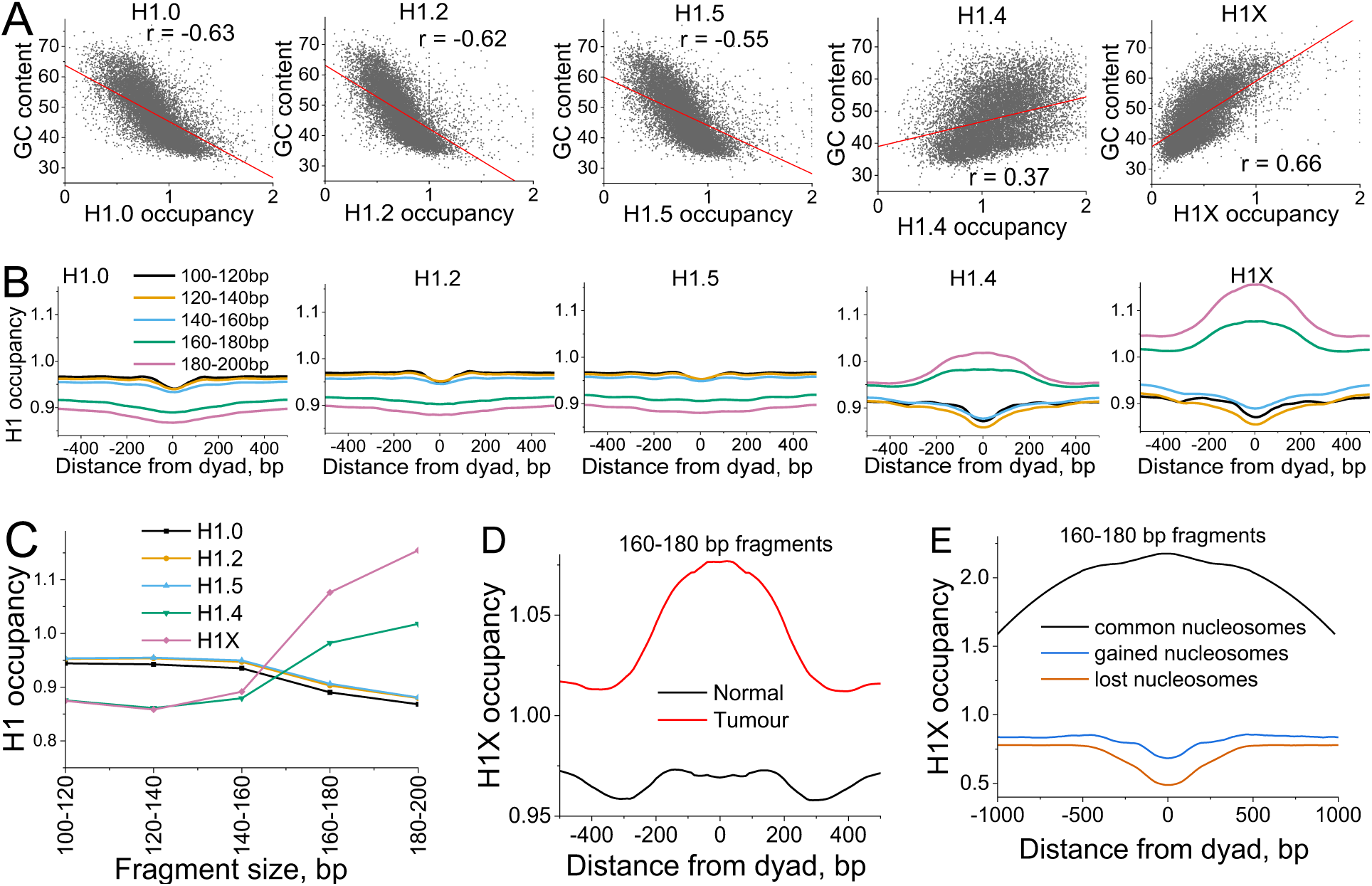
Effects of linker histones on nucleosome repositioning in breast cancer. A) Correlation of GC content and H1 occupancy in 10-kb regions genome-wide. H1 variants H1.0, H1.2 and H1.5 preferentially bind AT-rich regions, while H1.4 and H1X have a higher affinity for GC-rich regions. B-D) Averaged aggregate profiles of H1 occupancy around different types of DNA fragments protected from MNase digestion. These profiles were first calculated individually for each MNase-seq tumour tissue sample reported in this study, and then averaged across all patients. B) Occupancy of H1 variants around nucleosome dyads calculated for different DNA fragment sizes. C) H1 occupancy in the region [-50, 50] bp around nucleosome dyad calculated for different H1 variants as a function of DNA fragment size. D) Average H1X occupancy in T47D breast cancer cell line calculated around DNA fragments of 160-180 bp in normal (black lines) and tumour breast tissues (red). E) Average H1X occupancy profiles around the consensus set defined based on all tissue samples from this study, of common nucleosomes (black), and nucleosomes which are gained (blue) and lost in cancer (orange).

Given this relationship, the redistribution of H1.4 and H1X in BRC can lead to a significant depletion in the fraction of large-size nucleosomal DNA fragments, while the effect of H1.0, H1.2 and H1.5 histones is less pronounced (Figure 4C). A quantitatively similar effect was observed for nucleosome maps determined based on MNase-H3 (Supplementary Figure S7). Based on Figures 4A-C, linker histone H1X may have the largest effect on protection from the digestion of chromatosome particles (>160 bp). Therefore, we selected this histone variant for more advanced analysis. Figure 4D shows that genomic regions that give rise to protected DNA fragments with sizes 160-180 bp are characterised by a flat H1X profile in normal tissues but have a peak of H1X occupancy in the tumours. Furthermore, “common” 160-180 bp fragments protected from digestion both in normal and cancer tissues had ∼2-fold H1X enrichment, whereas regions that “lost” or “gained” nucleosomes in tumour had H1X depletion in BRC (Figure 4E). This suggests that H1X may be an important factor affecting the global nucleosome reorganisation in BRC.

### NRL shortening in breast cancer is not restricted to specific genomic regions

Genome- wide NRL changes could potentially arise due to cancer-specific changes in distinct types of genomic regions. To check this hypothesis, we calculated the NRL across all samples from our patient cohort in different types of genomic regions, such as active chromatin A-compartments (Figure 5A), regions surrounding gene promoters (Figure 5B), *Alu* repeats (Figure 5C), regions enriched with linker histone H1X (Figure 5D) and differentially methylated DNA regions (DMRs, Figure 5E). In line with previous studies (Valouev et al. 2011; Voong et al. 2016; Baldi et al. 2018a) and with Figure 2E, more active chromatin regions had a smaller NRL than less active regions. When comparing adjacent tumour versus non-tumour breast tissues for individual patients, all these types of genomic regions were characterised by an NRL decrease in tumour tissue. *Alu* repeats showed the largest heterogeneity among cancer patients, while DMRs showed less heterogeneity and one of the largest NRL changes (8 bp) between tumour and normal tissues. Regions enriched with H1X had a slightly smaller NRL change, followed by gene promoters and active A-compartments, consistent with the smaller cancer-specific NRL change inside the genes reported in Figure 2D. Thus, NRL shortening occurred in all studied types of genomic regions, although in a quantitatively different manner.

**Figure 5.**
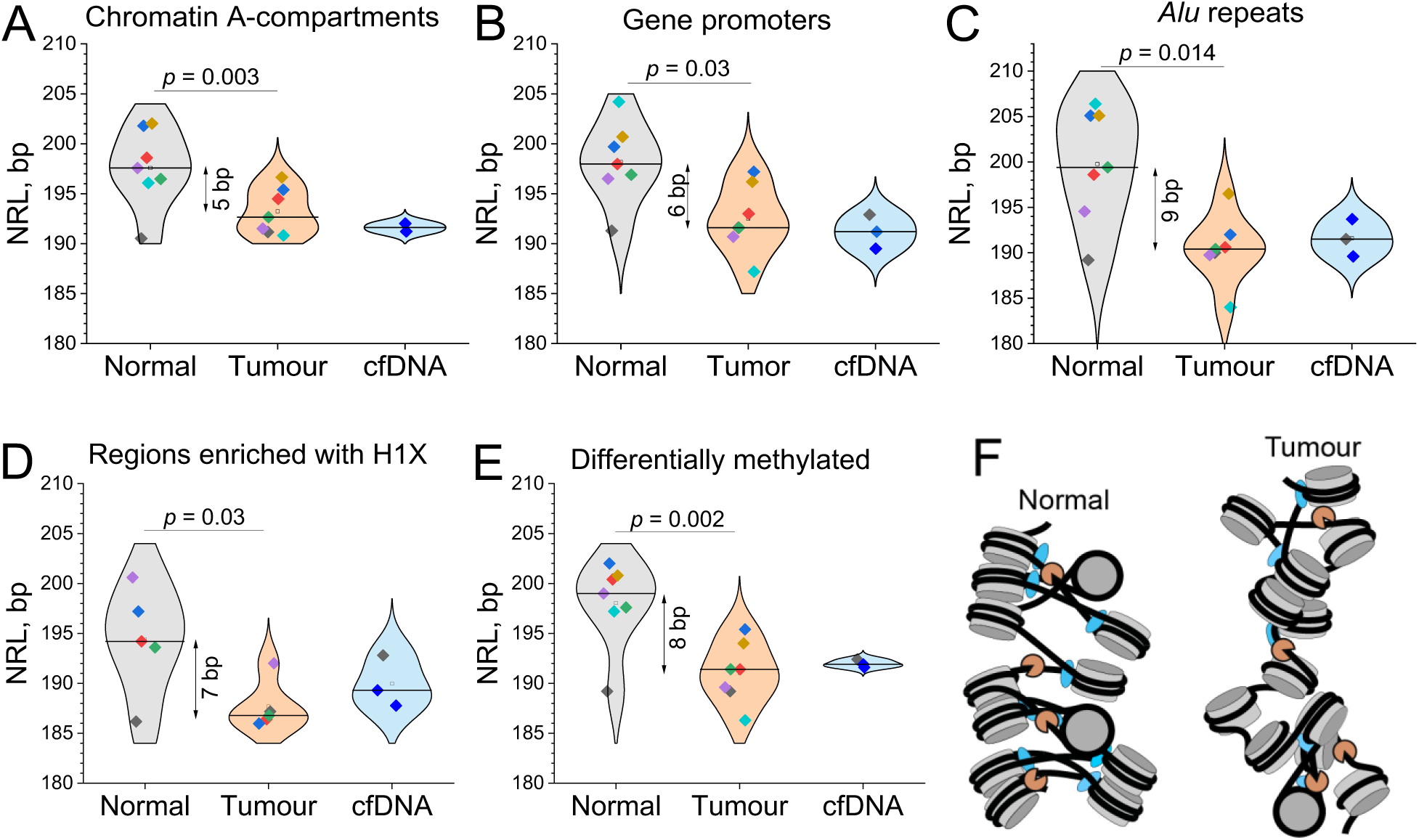
NRL shortening as a novel breast cancer marker. A-E) NRL values calculated for each individual sample from this study inside different types of genomic regions. A) Transcriptionally active A-compartments of chromatin annotated in breast cancer MCF-7 cells; B) Regions surrounding gene promoters. C) Genomic regions around Alu repeats; D) Regions enriched with H1X linker histones in breast cancer T47D cells; E) Differentially methylated DNA regions annotated based on cell lines MCF-10A, MCF-7 and MDA-MB-231. Open squares – average NRL values for a given condition; horizontal lines – median values. The colours of the points are assigned per sample as in Figure 3A. F) A scheme of the global change of nucleosome compaction in breast cancer tissues. DNA (black) is wrapped around histone octamers (grey) in contact with linker histones (blue). MNase used in this study or apoptotic nucleases present in situ (orange) cut DNA not protected by nucleosomes and linker histones. Cancer cells have shorter NRL and contain more nucleosomes which are less protected from MNase digestion due to redistribution of linker histone variant H1X, changes in DNA methylation and other factors.

## Discussion

Our studies have revealed systematic nucleosome changes in tumour cells that occur at two levels: (i) Small-scale changes at the level of individual nucleosomes that correspond to differential gene expression and TF binding (Figures 1 and 2); (ii) A genome-wide decrease in the NRL in tumour tissue (which holds true in all types of genomic regions that we analysed), as summarised in Figure 5F.

A small-scale BRC-associated nucleosome gain was particularly pronounced at CpG islands, promoters of genes involved in BRC signalling, and CTCF binding sites, while nucleosome loss was preferentially associated with L1 repeats and regulators of chromatin organisation. Such nucleosome repositioning was particularly pronounced at promoters of genes encoding DNA-binding proteins, which may be helpful to resolve one of the challenges in cfDNA-based nucleosomics, the determination of the tissue of origin (Snyder et al. 2016). In the current analysis we have focused instead on mechanistic effects, leaving the cfDNA tissue of origin problem for subsequent studies. Another noteworthy observation is that despite common assumptions in the field of cfDNA nucleosomics (Ulz et al. 2019; Doebley et al. 2022), not all TF binding sites are effective as cancer markers, even if they come from experimental binding maps (Figure 1D-F). For example, we found that only very stringent subsets of ERα sites allow one to stratify patients in terms of their differential nucleosome occupancy (Figure 1E).

Our finding that nucleosome reorganisation in cancer is characterised by a shorter NRL seemingly resembles the effect of shortening of cfDNA fragments in cancer patients in comparison with healthy people (Snyder et al. 2016; Underhill et al. 2016; Mouliere et al. 2018; Serpas et al. 2019; van der Pol and Mouliere 2019; Guo et al. 2020; Ding and Lo 2022). However, it is important to note that the NRL is more robust with respect to the experimental setup than the sizes of DNA fragments protected from MNase digestion (the former has been used historically as a biological characteristic of the cells, while the latter may depend on the level of MNase digestion). Indeed, we showed that even considering possible differences in chromatin digestion, the effect of NRL shortening in tumour tissue still stands (Figure 2C and F). While these effects can be uncoupled, their mechanisms may be related (e.g., through linker histone redistribution, changes of DNA methylation, etc).

Many classical studies were devoted to the investigation of NRL changes as a function of cell differentiation or cell state (e.g. see references in (Beshnova et al. 2014; Singh and Mueller-Planitz 2021)), and it is well established that NRL can change during differentiation. A hypothesis that the NRL can change in cancer was put forward initially in 1976 (Compton et al. 1976), and has gained traction again in recent publications, in particular relating this to linker histones H1 (Willcockson et al. 2021; Yusufova et al. 2021). To the best of our knowledge, we provide here the first evidence of a systematic NRL decrease in solid tumours based on adjacent tumour/non-tumour tissues from the same patients. Importantly, the effect of a global NRL decrease is different from the local effects of transcription-specific changes of nucleosome spacing as in Figure 2E at regulatory regions such as gene promoters (Valouev et al. 2011; Snyder et al. 2016) and TF binding sites (Teif et al. 2012; Teif et al. 2017; Clarkson et al. 2019). The global NRL decrease reported here for breast cancer may also take place in other cancers. Indeed, we have also found a similar genome-wide NRL shortening effect in B-cells from patients with chronic lymphocytic leukaemia, which was more pronounced for more aggressive cancer subtypes (Piroeva et al. 2023).

On a practical side, the finding that NRL decreases in tumour tissues may be directly relevant for the clinic, e.g., in relation to liquid biopsies based on cfDNA. Depending on the cancer stage, cfDNA in patients with solid tumours such as BRC may contain 1-30% of DNA from tumour tissues, while the majority of cfDNA usually comes from blood cells. If the nucleosome spacing changes characteristic for the tumour manifest in cfDNA, this may help to classify patients without the need to determine the cells of origin of cfDNA. The use of this effect in patient stratification is beyond the scope of the current publication but will be the subject of subsequent work. It is also worth noting that the small patient cohort considered here did not allow us to study systematically the effects of heterogeneity between patients. But even if the heterogeneity between patients is not known, this method may be still applied to monitoring a specific patient, with nucleosome spacing changes as a signal of cancer progression or response to therapy.

We showed here that the NRL decrease in BRC may be associated with several molecular mechanisms, including changes of gene expression, redistribution of linker histones, differential DNA methylation, changes in chromatin domains and the chromatin state of DNA sequence repeats. Since NRL shortening was not associated solely with any of these effects, it is likely that several mechanisms act synergistically. The clarification of the molecular mechanisms responsible for these effects will require a separate study, where DNA methylation and linker histones are measured in the same tissues for which the nucleosome maps have been determined. It is also intriguing that nucleosome-level changes may be connected to macroscopic effects, such as changes in the softness of cells/cell nuclei at the periphery of breast cancer tumours (Han et al. 2020), as well as changes in the physical properties of chromatin fibers as a feature of cancer progression (Stowers et al. 2019). It is worth noting that cancer cells have many stem cell-like properties. In particular, the chromatin of stem cells is characterised by smaller NRL (Teif et al. 2012) (Figure 2F). Thus, NRL shortening in tumour tissues may be related to de-differentiation of cancer cells (Friedmann-Morvinski and Verma 2014). On the mechanistic side, shorter NRL has been shown to change the topology of nucleosome fibres to more deformable and prone to macroscopic self-association (Zhurkin and Norouzi 2021), consistent with Figure 5F. Thus the 5-10 bp decrease in NRL in breast tissues may be associated with dramatic genome reorganisation, causative mechanistic details of which remain unknown.

Taken together, this study provides the first experimental nucleosome maps in paired tumour/normal breast tissues, which will be a valuable resource for the community, and reported conceptually new effects that deserve subsequent studies analysing the corresponding mechanisms. Furthermore, these new effects allow us to establish some of the missing theoretical foundations for cfDNA-based nucleosomics analysis. These effects are probably not limited to BRC, and future studies are needed to expand this methodology to larger patient cohorts and more cancer types.

## Methods

### Human tissues and blood

We studied matched tissue samples and blood samples from the same individual patients as described below. Primary tumour tissues together with paired peripheral tissues (also called “normal”) were collected during surgery from breast cancer patients numbered P1-P4 treated at the Colchester General Hospital. Tumour tissues were taken from the center of the tumour, and normal tissues were taken from a distance of about 1 millimetre from the periphery. For patients P1 and P3 in addition matching blood samples were drawn before the surgical procedure. Samples were stored at -80 °C. Detailed morphological and molecular characterisation of the paired normal/tumour breast tissues from the patient cohort to which the patients studied here belong has been performed in our previous publications (Docquier et al. 2005; D’Arcy et al. 2008; Docquier et al. 2009). The paired peripheral tissues displayed morphological characteristics identical to normal breast tissues whereby the following structures were present: breast ducts and lobules with two cell layers consisting of inner luminal/epithelial cells and outer basal/myoepithelial cells, blood vessels and stromal cells. Two tissue types had distinct proliferative states. Normal breast tissues were negative for the proliferation marker Ki-67, whereas breast tumour tissues were positive (see Supplemental Figure 4 in Docquier et al., 2009 (Docquier et al. 2009)). The cell composition of the paired breast tissues from this cohort was also clarified in our previous work (Docquier et al. 2009) using immunofluorescent staining with markers specific for different cell types: CK14 for myoepithelial cells and CK19 for luminal cells. Normal breast tissues were composed of myoepithelial cells and luminal cells. Tumour tissues were composed predominantly of luminal cells. Detailed clinical information for patients P1 and P2 provided by the Colchester General Hospital is available in Supplementary Table S1. Both patients P1 and P2 are ER-positive. For patient P3, we determined ERα status using Western blot (Supplementary Figure S1), which did not detect ERα presence in the tumour sample. The study was approved by the local ethics committee (NRES Committee East of England, reference number MH363).

### MNase-seq and MNase-assisted histone H3 ChIP-seq (MNase-H3)

#### Chromatin extraction

Frozen tissue was first ground into fine powder in liquid nitrogen. It was further homogenized by manual douncing in 5 ml ice-cold Buffer I (0.3 M Sucrose in 60 mM KCl, 15 mM NaCl, 5mM MgCl_2_, 0.1 mM EGTA, 15 mM Tris-HCl (pH 7.5) with freshly added 0.5 mM DTT, 0.1 mM PMSF and 1XProtease Inhibitor Cocktail, Thermo Scientific Halt Protease Inhibitor Single-use Cocktail (100X). The homogenized cell suspension was centrifuged in a 15 ml Falcon conical centrifuge tube at 6000 g for 10 minutes at 4 °C. The precipitated cells were re-suspended in 500 µL of Buffer I, then 500 µL of Buffer II (same as Buffer I, but with freshly added 0.4 % (v/v) IGEPAL CA-630) was gently added on top and the lysate was incubated on ice for 3 minutes, followed by centrifugation at 3000 g for 3 minutes at 4 °C. The pellet was re-suspended in 1 ml of ice-cold NBR Buffer (85 mM NaCl, 5.5 % Sucrose, 10 mM Tris-HCl pH 7.5, 3 mM MgCl_2,_ 1.5 mM CaCl_2_ with freshly added 0.2 mM phenylmethylsulfonyl fluoride (PMSF) and 1 mM DTT) and centrifuged at 3000 g for 3 minutes at 4 °C. The pellet then was re-suspended in 200 µl of NBR buffer. Samples were treated with 1 µl RNAse A (10 mg/ml) per 100 µl of nuclear lysate for 5 minutes at room temperature. *MNase digestion*. MNase restriction enzyme (Sigma, Cat No: N5386-500UN) was diluted to 1U (Sigma Boehringer Units) in MNase buffer (50 % Glycerol, 10 mM Tris pH7.6 and 50 mM NaCl). The pellets re-suspended in 200 µl of NBR buffer in the chromatin extraction step were digested with MNase enzyme as follows. Chromatin derived from paired normal and tumour tissue was diluted with NBR buffer to keep the same concentration in both preparations. A 20 µl aliquot of purified nuclei was test digested with 1 µl MNase for 7 min at room temperature to assess the optimal digestion time and enzyme concentrations. 1 µg of chromatin was digested with 1U of MNase for 7 minutes at room temperature to achieve 80:20 relative abundance of mono- and di-nucleosome fractions on the agarose gel. The MNase digestion was stopped by addition of equal volume (NBR:STOP, 1:1) of STOP Buffer (215 mM NaCl, 10 mM TrisHCl pH 8, 20 mM EDTA, 5.5 % Sucrose, 2 % TritonX 100 with freshly added 0.2 mM PMSF, 1 mM DTT and 2X Protease Inhibitors). After digestion, chromatin was incubated on ice in the fridge at 4 °C overnight to release the digested chromatin.

#### Chromatin Immunoprecipitation (ChIP)

10 µl of Protein A beads (Dynabeads^TM^ Protein A; Invitrogen) equivalent to 10 µl bed volume of slurry were prepared per each ChIP reaction. Bead storage buffer was aspirated, and the beads were washed twice in 500 µl in Block solution (0.5 % BSA (w/v) in PBS supplemented with 0.1 mM PMSF). Beads sufficient for 4 ChIP samples were all combined and dissolved in 600 µl of Block solution with the addition of 5 µg of anti-H3 antibody Abcam AB1791 per ChIP reaction (20 µg for 40 µl of beads for 4 ChIP samples). The antibody-bead solution was incubated on a slowly rotating wheel at 4 °C for two hours. After incubation, the beads were again washed in 600 µl block solution to remove unbound antibodies. The chromatin was incubated overnight after MNase digestion and then centrifuged for 10 minutes at a max speed of 9800 g at 4 °C. The supernatant with released chromatin was collected and stored at -20 °C and used for the ChIP experiment. 5 % of chromatin volume used for MNase-assisted ChIP-seq was frozen at -20 °C as Input and used for MNase-seq. 300 µl of chromatin was incubated with the antibody-bound Protein A beads for 3 hours on a rotating wheel at 4 °C. Unbound chromatin was washed 4 times in 1 ml volume of ChIP-Wash buffer (150 mM NaCl, 10 mM Tris-HCl pH 8, 2 mM EDTA, 1 % NP40, 1 % Nadeoxycholate with freshly added 0.5 mM DTT, 0.1 mM PMSF and 1XProtease Inhibitor Cocktail). The initial wash of bound beads was done by gently mixing beads in the ChIP-Wash buffer for one min. For the subsequent 3 washes: the tube was rotated for 5 minutes at 4 °C on a rotating wheel, then placed in a magnetic rack for 1 min at 4 °C and then the supernatant was withdrawn. After two washes the beads with wash buffer were transferred to new tubes. And as a final step, the beads were washed with 1 ml of filtered 1x TE buffer (10 mM Tris-HCl containing 1 mM EDTA) at room temperature, after which the TE buffer was removed as above. The elution of bound chromatin was performed with 100 µl of Elution buffer (0.1 M NaHCO_3,_ 1 % SDS) supplemented with 2 µl of Proteinase K per ChIP sample tube. The beads were incubated with the elution buffer for 6-12 hours at 55 °C. Similarly, 2 µl of Proteinase K were added to the Input material and incubated at 55 °C for 6-12 hours. The incubated beads were briefly vortexed and the supernatant was collected by placing the tubes in a magnetic rack. The beads were additionally washed in 20 µl of elution buffer and the supernatant collected as above. The eluants were further purified with PCR purification columns according to the manufacturer’s instructions and eluted in 20 µl of elution buffer of the kit. These final eluants were stored at -20 °C ready for the library preparation step.

### Cell culture

A549 and T47D cells were acquired from the ATCC and cultured in Dulbecco’s Modified Eagle’s Medium (DMEM, Corning, NY, USA) or Roswell Park Memorial Institute (RPMI, Corning) medium 1640, respectively. Media were supplemented with 10% FCS and 2 mM L-glutamine, 100 U/ml penicillin, 100 mg/ml streptomycin (PSG, Sigma Aldrich, MI, USA) and cultured as previously described (Brooke et al. 2014).

### Immunoblotting

Samples were lysed in RIPA buffer (150 mM NaCl, 1% IGEPAL CA-630, 0.5% sodium deoxycholate, 0.1% SDS, 25 mM Tris pH 7.4) containing freshly added protease inhibitors (Halt Protease Cocktail, Thermo Fisher, Waltham, MA, USA) and prepared as previously described (Leach et al. 2021). The DC protein assay (BioRad, Hercules, CA, USA) was used to quantify protein concentrations. 60 μg of protein was separated using SDS-PAGE and immunoblotting performed. Primary antibodies used were rabbit anti-ERα (1:1000, sc- 543, Santa Cruz, TX, USA), mouse α-tubulin (1:10,000, B-5-1-2, Sigma Aldrich). Secondary HRP-conjugated antibodies (Sigma Aldrich) were used at 1:2000. Proteins were visualised using Immobilon Forte HRP substrate (Merck Millipore, MA, USA) and a Fusion FX Chemi Imager (Vilber Lourmat, Collégien, France).

### External datasets

Raw data from ChIP-seq of histone H1 variants in T47D cells was obtained from GEO entry GSE156036 (Serna-Pujol et al. 2022). Coordinates of chromatin A- and B-compartments defined based on Hi-C in MCF-7 cells (Achinger-Kawecka et al. 2020) were kindly provided by the authors in human genome assembly hg38 and converted to hg19 using LiftOver utility of the UCSC Genome Browser. Coordinates of genomic regions with differential DNA methylation between breast cell lines MCF-10A, MCF-7 and MDA-MB-231 were obtained from (Lee et al. 2020). Whole-genome bisulfite sequencing in MCF-7 cells reported by (Lyu et al. 2021) was obtained from GSM4055031 in the form of methylation calls in hg19 genome assembly. Coordinates of chromatin domains in breast cells enriched with H3K9me2/3 determined with ChIP-seq by the ENCODE consortium (ENCODE 2012) were obtained from the following GEO entries: H3K9me3 in HMEC cells (GSM1003485), H3K9me3 in MCF-7 cells (GSE96517), H3K9me2 in MCF-7 cells (GSE96141). ERα binding sites were defined based on ChIP-seq peaks in six female BRC patients reported by Severson et al (Severson et al. 2018) (GSE104399). We constructed two consensus datasets of ERα binding: more stringent dataset contained the overlapping peaks appearing in all six patients (193 peaks), and less stringent dataset of peaks confirmed in any three out of these six patients (11,637 peaks). These datasets were further refined with RSAT (Santana-Garcia et al. 2022) to consider only ERα motifs inside these peaks. FOSL2 binding sites were defined by re-calling peaks using HOMER starting from raw ChIP-seq data in MCF-7 cells reported by the ENCODE consortium (GSM1010768) (ENCODE 2012). This resulted in 14,106 peaks, which were then narrowed down using the coordinates of FOSL2 motifs inside these peaks determined with RSAT’s default settings. CTCF binding sites in MCF-7 reported by the ENCODE consortium (ENCODE 2012) were obtained from GSM822305. Processed gene expression in breast invasive carcinoma reported by the TCGA Consortium (Hoadley et al. 2018) was obtained from cBioPortal (www.cbioportal.org) and sorted based on RSEM values. Top 3,000 (highest expression) versus bottom 3,000 genes (lowest expression) were selected from this dataset based on the RSEM values averaged across either normal or cancer samples and used in the analysis in Figure 2E. MNase-seq in MCF-7 was obtained from GSE77526 and GSE51097 (Shimbo et al. 2013). The latter dataset contains separate samples for eight different levels of MNase digestions, which we have processed separately. MNase-seq in T47D cells was obtained from GSE74308 (Lavender et al. 2016); MNase-seq in human embryonic stem cells (hESC) from GSE49140 (Yazdi et al. 2015). Unless specified otherwise, we mapped each sample using Bowtie2 to hg19, considering only uniquely mapped reads with up to 1 mismatch.

### Calculation of aggregate occupancy profiles

Aggregate occupancy profiles and nucleotide frequency profiles were calculated separately for each sample using HOMER (Heinz et al. 2010). Where aggregate profiles are reported averaged per condition and per experiment type, the HOMER-generated profiles, each normalised individually to the sequencing depths of the corresponding sample, were plotted in Origin Pro (originlab.com) and the arithmetic average was determined across all samples with the same condition and the same experiment type. H1 occupancy profiles were normalised by dividing the ChIP-seq signal by the corresponding Input.

### NRL calculation

The method of NRL calculation from paired-end reads was developed in our previous publication (Teif et al. 2012) and is available in NucTools software package (Vainshtein et al. 2017). This method is conceptually similar to the method based on single-end reads (Valouev et al. 2011) but offers higher precision because it’s based directly on nucleosome centers (dyads) and applies different filtering. The NucTools protocol is based on the calculation of the distribution of frequencies of distances between nucleosome dyads with single-bp resolution, separately per each chromosome and then averaged over all chromosomes. At the next step, Origin Pro (originlabs.com) and NRLcalc package (Clarkson et al. 2019) were used to detect peaks of the sinusoidal oscillations of the distribution of frequencies of distances between nucleosome dyads, followed by linear regression to determine the NRL. The calculations performed here were done with NucTools using up to 40 million reads per chromosome, considering dyad-dyad distances up to 2,000 bp, and discarding cases with more than 50 mapped reads centred on the same base pair. Changing the cutoff to 20, 30 and 40 reads per base pair did not affect the NRL values. NRLs were calculated separately for each sample, both genome-wide and inside sets of genomic regions of interest. For the calculation of NRL inside genomic regions, we defined them as follows: regions surrounding promoters were defined as intervals [-5,000; 5,000] from annotated RefSeq TSSs. The genes used in NRL calculation were selected as protein-coding genes according to the UCSC Genome Browser annotation, with the length satisfying criteria 1 kb < Length < 200 kb. This resulted in 16,161 genes. For the calculations in Figures 2E and F these genes were further classified as top 3,000 genes which are highly- and lowly-expressed based on TCGA cancer genome atlas RSEM expression values (Hoadley et al. 2018), as well as top 3,000 genes enriched with MNase-seq fragments of sizes in one of the following ranges [100-120 bp], [120-140 bp], [140-160 bp], [160-180 bp], [180-200 bp]. Alu repeats annotated based on UCSC Genome Browser’s Repeat Masker were extended by 1,000 bp on both sides. DMR regions defined above (Lee et al. 2020) were also extended by 1,000 bp on both sides. Regions enriched with H1X were defined based on MACS2 peak calling with default parameters using ChIP-seq of H1X in the T47D breast cancer cell line reported in (Serna-Pujol et al. 2022), resulting in 33,337 peaks which were extended by 1,000 bp on both sides. Genes enriched and depleted with protected DNA fragments of size 160-180 bp used in Figure 2C were defined by calculating the number of 160-180 bp fragments per gene normalised by the gene length, then sorting the genes based on this number and defining the top 3,000 genes as enriched with “long nucleosomes” and the bottom 3,000 genes as depleted of “long nucleosomes”. Regions with low- and high-GC content used in Figure 2D were defined by calculating average GC content with the running 10 kb window and defining equal numbers of low-GC and high-GC regions (144,758 regions of each type, threshold GC content 40%). Genes marked by differential nucleosome occupancy in their promoters used in Figure 2D were defined by intersecting the genes from the set 1 kb < Length < 200 kb defined above with the sets of nucleosomes gained and lost in patient P2 in BRC versus normal tissue (Table S2). This resulted in 1,423 genes that gained nucleosomes at their promoters and 401 genes lost nucleosomes at their promoters.

### Calculation of aggregate ChIP-seq occupancy profiles

Aggregate occupancy profiles and nucleotide frequency profiles were calculated separately for each sample using HOMER (Heinz et al. 2010) and NucTools (Vainshtein et al. 2017). Where aggregate profiles are reported averaged per condition and per experiment type, the HOMER-generated profiles, each normalised individually to the sequencing depths of the corresponding sample, were plotted together in Origin Pro (originlab.com) and the arithmetic average was determined across all samples with the same condition and the same experiment type. H1 occupancy profiles were normalised by dividing the ChIP-seq signal by the corresponding Input.

### Calculation of aggregate DNA methylation profiles

We used bisulfite sequencing data in MCF-7 cells (Lyu et al. 2021), with methylation beta-values determined by the authors. NucTools (Vainshtein et al. 2017) was used to split the regions into chromosomes and 5mC occurrence per CpG was calculated with a cfDNAtools script bed2occupancy.v3d.methyl.pl as in (Piroeva et al. 2023). Aggregate DNA methylation profiles were calculated around centres of common and shifted nucleosomes defined above using NucTools, summing up the methylation beta-values for each CpG located at a given distance from the genomic feature of interest as in (Wiehle et al. 2019). When the corresponding graph refers to “DNA methylation (a.u.)”, this corresponds to the value obtained by summation of all corresponding beta-values without further normalization.

### Enrichment analysis

BedTools “Fisher” function was used to calculate fold enrichment as well as the corresponding *P*-values of the Fisher test for the enrichment with respect to random genomic regions for gained and lost nucleosomes inside gene promoters (defined as +/-1000 bp from RefSeq-annotated transcription start sites (TSSs)); CpG islands, Alu repeats, L1 repeats, and microsatellite repeats based on the UCSC Genome Browser Repeat Masker; CTCF sites defined using ChIP-seq by the ENCODE consortium (ENCODE 2012); DMRs (Lee et al. 2020) and H1.4 and H1X (Serna-Pujol et al. 2022) defined as detailed above.

### Gene annotation

RefSeq-annotated gene promoters defined as +/-1000 bp from TSS were intersected with regions with “lost” and “gained” nucleosomes defined above, considering all fragment sizes between 120-180 bp. Gene Ontology analysis was performed using the R package gprofiler2 (Kolberg et al. 2020). Gene network was constructed with Cytoscape 3.9.1 (Shannon et al. 2003) using STRING database (Szklarczyk et al. 2023), including genes with promoters marked by nucleosome gain in BRC, associated with gene ontology terms related to “DNA-binding”.

## Data availability

The processed sequencing data generated in this study have been submitted to the NCBI Gene Expression Omnibus (GEO) and will be made available upon journal publication.

## Acknowledgements

We thank Fred Winston (Harvard Medical School) for valuable comments on the manuscript and Stuart Newman (University of Essex) for the computer cluster support.

## References

Achinger-Kawecka J, Valdes-Mora F, Luu P-L, Giles KA, Caldon CE, Qu W, Nair S, Soto S, Locke WJ, Yeo-Teh NS. 2020. Epigenetic reprogramming at estrogen-receptor binding sites alters 3D chromatin landscape in endocrine-resistant breast cancer. Nature communications 11: 1–17.

Baldi S, Jain DS, Harpprecht L, Zabel A, Scheibe M, Butter F, Straub T, Becker PB. 2018a. Genome-wide Rules of Nucleosome Phasing in Drosophila. Mol Cell 72: 661–672 e664.

Baldi S, Krebs S, Blum H, Becker PB. 2018b. Genome-wide measurement of local nucleosome array regularity and spacing by nanopore sequencing. Nat Struct Mol Biol 25: 894–901.

Bansal N, Petrie K, Christova R, Chung CY, Leibovitch BA, Howell L, Gil V, Sbirkov Y, Lee E, Wexler J et al. 2015. Targeting the SIN3A-PF1 interaction inhibits epithelial to mesenchymal transition and maintenance of a stem cell phenotype in triple negative breast cancer. Oncotarget 6: 34087–34105.

Beshnova DA, Cherstvy AG, Vainshtein Y, Teif VB. 2014. Regulation of the nucleosome repeat length in vivo by the DNA sequence, protein concentrations and long-range interactions. PLoS Comput Biol 10: e1003698.

Brett JO, Spring LM, Bardia A, Wander SA. 2021. ESR1 mutation as an emerging clinical biomarker in metastatic hormone receptor-positive breast cancer. Breast Cancer Research 23: 85.

Brooke GN, Powell SM, Lavery DN, Waxman J, Buluwela L, Ali S, Bevan CL. 2014. Engineered repressors are potent inhibitors of androgen receptor activity. Oncotarget 5: 959–969.

Chereji RV, Kan TW, Grudniewska MK, Romashchenko AV, Berezikov E, Zhimulev IF, Guryev V, Morozov AV, Moshkin YM. 2016. Genome-wide profiling of nucleosome sensitivity and chromatin accessibility in Drosophila melanogaster. Nucleic Acids Res 44: 1036–1051.

Clarkson CT, Deeks EA, Samarista R, Mamayusupova H, Zhurkin VB, Teif VB. 2019. CTCF-dependent chromatin boundaries formed by asymmetric nucleosome arrays with decreased linker length. Nucleic Acids Res 47: 11181–11196.

Cole L, Kurscheid S, Nekrasov M, Domaschenz R, Vera DL, Dennis JH, Tremethick DJ. 2021. Multiple roles of H2A.Z in regulating promoter chromatin architecture in human cells. Nat Commun 12: 2524.

Compton JL, Bellard M, Chambon P. 1976. Biochemical evidence of variability in the DNA repeat length in the chromatin of higher eukaryotes. Proc Natl Acad Sci U S A 73: 4382–4386.

D’Arcy V, Pore N, Docquier F, Abdullaev ZK, Chernukhin I, Kita GX, Rai S, Smart M, Farrar D, Pack S et al. 2008. BORIS, a paralogue of the transcription factor, CTCF, is aberrantly expressed in breast tumours. British Journal of Cancer 98: 571–579.

Ding SC, Lo YMD. 2022. Cell-Free DNA Fragmentomics in Liquid Biopsy. Diagnostics (Basel) 12.

Docquier F, Farrar D, D’Arcy V, Chernukhin I, Robinson AF, Loukinov D, Vatolin S, Pack S, Mackay A, Harris RA et al. 2005. Heightened Expression of CTCF in Breast Cancer Cells Is Associated with Resistance to Apoptosis. Cancer Research 65: 5112–5122.

Docquier F, Kita G-X, Farrar D, Jat P, O’Hare M, Chernukhin I, Gretton S, Mandal A, Alldridge L, Klenova E. 2009. Decreased Poly(ADP-Ribosyl)ation of CTCF, a Transcription Factor, Is Associated with Breast Cancer Phenotype and Cell Proliferation. Clinical Cancer Research 15: 5762–5771.

Doebley AL, Ko M, Liao H, Cruikshank AE, Santos K, Kikawa C, Hiatt JB, Patton RD, De Sarkar N, Collier KA et al. 2022. A framework for clinical cancer subtyping from nucleosome profiling of cell-free DNA. Nat Commun 13: 7475.

ENCODE. 2012. An integrated encyclopedia of DNA elements in the human genome. Nature 489: 57–74.

Friedmann-Morvinski D, Verma IM. 2014. Dedifferentiation and reprogramming: origins of cancer stem cells. EMBO Rep 15: 244–253.

Guo J, Ma K, Bao H, Ma X, Xu Y, Wu X, Shao YW, Jiang M, Huang J. 2020. Quantitative characterization of tumor cell-free DNA shortening. BMC Genomics 21: 473.

Han YL, Pegoraro AF, Li H, Li K, Yuan Y, Xu G, Gu Z, Sun J, Hao Y, Gupta SK et al. 2020. Cell swelling, softening and invasion in a three-dimensional breast cancer model. Nat Phys 16: 101–108.

Heinz S, Benner C, Spann N, Bertolino E, Lin YC, Laslo P, Cheng JX, Murre C, Singh H, Glass CK. 2010. Simple combinations of lineage-determining transcription factors prime cis-regulatory elements required for macrophage and B cell identities. Mol Cell 38: 576–589.

Hoadley KA, Yau C, Hinoue T, Wolf DM, Lazar AJ, Drill E, Shen R, Taylor AM, Cherniack AD, Thorsson V et al. 2018. Cell-of-Origin Patterns Dominate the Molecular Classification of 10,000 Tumors from 33 Types of Cancer. Cell 173: 291–304 e296.

Kolberg L, Raudvere U, Kuzmin I, Vilo J, Peterson H. 2020. gprofiler2 -- an R package for gene list functional enrichment analysis and namespace conversion toolset g:Profiler. F1000Res 9.

Lavender CA, Cannady KR, Hoffman JA, Trotter KW, Gilchrist DA, Bennett BD, Burkholder AB, Burd CJ, Fargo DC, Archer TK. 2016. Downstream Antisense Transcription Predicts Genomic Features That Define the Specific Chromatin Environment at Mammalian Promoters. PLoS genetics 12: e1006224.

Leach DA, Mohr A, Giotis ES, Cil E, Isac AM, Yates LL, Barclay WS, Zwacka RM, Bevan CL, Brooke GN. 2021. The antiandrogen enzalutamide downregulates TMPRSS2 and reduces cellular entry of SARS-CoV-2 in human lung cells. Nat Commun 12: 4068.

Lee I, Razaghi R, Gilpatrick T, Molnar M, Gershman A, Sadowski N, Sedlazeck FJ, Hansen KD, Simpson JT, Timp W. 2020. Simultaneous profiling of chromatin accessibility and methylation on human cell lines with nanopore sequencing. Nature Methods 17: 1191–1199.

Lidor Nili E, Field Y, Lubling Y, Widom J, Oren M, Segal E. 2010. p53 binds preferentially to genomic regions with high DNA-encoded nucleosome occupancy. Genome Res 20: 1361–1368.

Lyu R, Zhu X, Shen Y, Xiong L, Liu L, Liu H, Wu F, Argueta C, Tan L. 2021. Tumour suppressor TET2 safeguards enhancers from aberrant DNA methylation and epigenetic reprogramming in ERα-positive breast cancer cells. Epigenetics: 1–15.

Mouliere F, Chandrananda D, Piskorz AM, Moore EK, Morris J, Ahlborn LB, Mair R, Goranova T, Marass F, Heider K et al. 2018. Enhanced detection of circulating tumor DNA by fragment size analysis. Sci Transl Med 10.

Piroeva KV, McDonald C, Xanthopoulos C, Fox C, Clarkson CT, Mallm J-P, Vainshtein Y, Ruje L, Klett LC, Stilgenbauer S et al. 2023. Nucleosome repositioning in chronic lymphocytic leukaemia. Genome Res doi:10.1101/gr.277298.122: Advanced access.

Santana-Garcia W, Castro-Mondragon JA, Padilla-Galvez M, Nguyen NTT, Elizondo-Salas A, Ksouri N, Gerbes F, Thieffry D, Vincens P, Contreras-Moreira B et al. 2022. RSAT 2022: regulatory sequence analysis tools. Nucleic Acids Res 50: W670–W676.

Serna-Pujol N, Salinas-Pena M, Mugianesi F, Le Dily F, Marti-Renom MA, Jordan A. 2022. Coordinated changes in gene expression, H1 variant distribution and genome 3D conformation in response to H1 depletion. Nucleic Acids Res 50: 3892–3910.

Serpas L, Chan RWY, Jiang P, Ni M, Sun K, Rashidfarrokhi A, Soni C, Sisirak V, Lee WS, Cheng SH et al. 2019. Dnase1l3 deletion causes aberrations in length and end-motif frequencies in plasma DNA. Proc Natl Acad Sci U S A 116: 641–649.

Severson TM, Kim Y, Joosten SEP, Schuurman K, van der Groep P, Moelans CB, Ter Hoeve ND, Manson QF, Martens JW, van Deurzen CHM et al. 2018. Characterizing steroid hormone receptor chromatin binding landscapes in male and female breast cancer. Nat Commun 9: 482.

Shannon P, Markiel A, Ozier O, Baliga NS, Wang JT, Ramage D, Amin N, Schwikowski B, Ideker T. 2003. Cytoscape: a software environment for integrated models of biomolecular interaction networks. Genome Res 13: 2498–2504.

Shimbo T, Du Y, Grimm SA, Dhasarathy A, Mav D, Shah RR, Shi H, Wade PA. 2013. MBD3 localizes at promoters, gene bodies and enhancers of active genes. PLoS genetics 9: e1004028.

Shtumpf M, Piroeva KV, Agrawal SP, Jacob DR, Teif VB. 2022. NucPosDB: a database of nucleosome positioning in vivo and nucleosomics of cell-free DNA. Chromosoma 131: 19–28.

Singh AK, Mueller-Planitz F. 2021. Nucleosome Positioning and Spacing: From Mechanism to Function. J Mol Biol 433: 166847.

Snyder MW, Kircher M, Hill AJ, Daza RM, Shendure J. 2016. Cell-free DNA Comprises an In Vivo Nucleosome Footprint that Informs Its Tissues-Of-Origin. Cell 164: 57–68.

Stowers RS, Shcherbina A, Israeli J, Gruber JJ, Chang J, Nam S, Rabiee A, Teruel MN, Snyder MP, Kundaje A et al. 2019. Matrix stiffness induces a tumorigenic phenotype in mammary epithelium through changes in chromatin accessibility. Nat Biomed Eng 3: 1009–1019.

Szklarczyk D, Kirsch R, Koutrouli M, Nastou K, Mehryary F, Hachilif R, Gable AL, Fang T, Doncheva NT, Pyysalo S et al. 2023. The STRING database in 2023: protein-protein association networks and functional enrichment analyses for any sequenced genome of interest. Nucleic Acids Res 51: D638–D646.

Takaku M, Grimm SA, Shimbo T, Perera L, Menafra R, Stunnenberg HG, Archer TK, Machida S, Kurumizaka H, Wade PA. 2016. GATA3-dependent cellular reprogramming requires activation-domain dependent recruitment of a chromatin remodeler. Genome Biol 17: 36.

Teif VB, Beshnova DA, Vainshtein Y, Marth C, Mallm JP, Hofer T, Rippe K. 2014. Nucleosome repositioning links DNA (de)methylation and differential CTCF binding during stem cell development. Genome Res 24: 1285–1295.

Teif VB, Gould TJ, Clarkson CT, Boyd L, Antwi EB, Ishaque N, Olins AL, Olins DE. 2020. Linker histone epitopes are hidden by in situ higher-order chromatin structure. Epigenetics Chromatin 13: 26.

Teif VB, Mallm JP, Sharma T, Mark Welch DB, Rippe K, Eils R, Langowski J, Olins AL, Olins DE. 2017. Nucleosome repositioning during differentiation of a human myeloid leukemia cell line. Nucleus 8: 188–204.

Teif VB, Vainshtein Y, Caudron-Herger M, Mallm JP, Marth C, Hofer T, Rippe K. 2012. Genome-wide nucleosome positioning during embryonic stem cell development. Nat Struct Mol Biol 19: 1185–1192.

Ulz P, Perakis S, Zhou Q, Moser T, Belic J, Lazzeri I, Wolfler A, Zebisch A, Gerger A, Pristauz G et al. 2019. Inference of transcription factor binding from cell-free DNA enables tumor subtype prediction and early detection. Nat Commun 10: 4666.

Underhill HR, Kitzman JO, Hellwig S, Welker NC, Daza R, Baker DN, Gligorich KM, Rostomily RC, Bronner MP, Shendure J. 2016. Fragment Length of Circulating Tumor DNA. PLoS genetics 12: e1006162.

Vainshtein Y, Rippe K, Teif VB. 2017. NucTools: analysis of chromatin feature occupancy profiles from high-throughput sequencing data. BMC Genomics 18: 158.

Valouev A, Johnson SM, Boyd SD, Smith CL, Fire AZ, Sidow A. 2011. Determinants of nucleosome organization in primary human cells. Nature 474: 516–520.

van der Pol Y, Mouliere F. 2019. Toward the Early Detection of Cancer by Decoding the Epigenetic and Environmental Fingerprints of Cell-Free DNA. Cancer Cell 36: 350–368.

Voong LN, Xi L, Sebeson AC, Xiong B, Wang JP, Wang X. 2016. Insights into Nucleosome Organization in Mouse Embryonic Stem Cells through Chemical Mapping. Cell 167: 1555–1570 e1515.

Wiehle L, Thorn GJ, Raddatz G, Clarkson CT, Rippe K, Lyko F, Breiling A, Teif VB. 2019. DNA (de)methylation in embryonic stem cells controls CTCF-dependent chromatin boundaries. Genome research 29: 750–761.

Willcockson MA, Healton SE, Weiss CN, Bartholdy BA, Botbol Y, Mishra LN, Sidhwani DS, Wilson TJ, Pinto HB, Maron MI et al. 2021. H1 histones control the epigenetic landscape by local chromatin compaction. Nature 589: 293–298.

Yang X, Cai G-X, Han B-W, Guo Z-W, Wu Y-S, Lyu X, Huang L-M, Zhang Y-B, Li X, Ye G-L et al. 2021. Association between the nucleosome footprint of plasma DNA and neoadjuvant chemotherapy response for breast cancer. npj Breast Cancer 7: 35.

Yazdi PG, Pedersen BA, Taylor JF, Khattab OS, Chen YH, Chen Y, Jacobsen SE, Wang PH. 2015. Nucleosome Organization in Human Embryonic Stem Cells. PLoS One 10: e0136314.

Yusufova N, Kloetgen A, Teater M, Osunsade A, Camarillo JM, Chin CR, Doane AS, Venters BJ, Portillo-Ledesma S, Conway J et al. 2021. Histone H1 loss drives lymphoma by disrupting 3D chromatin architecture. Nature 589: 299–305.

Zhurkin VB, Norouzi D. 2021. Topological polymorphism of nucleosome fibers and folding of chromatin. Biophys J 120: 577–585.

