## Supplementary Materials for "Nucleosome reorganisation in breast cancer tissues"

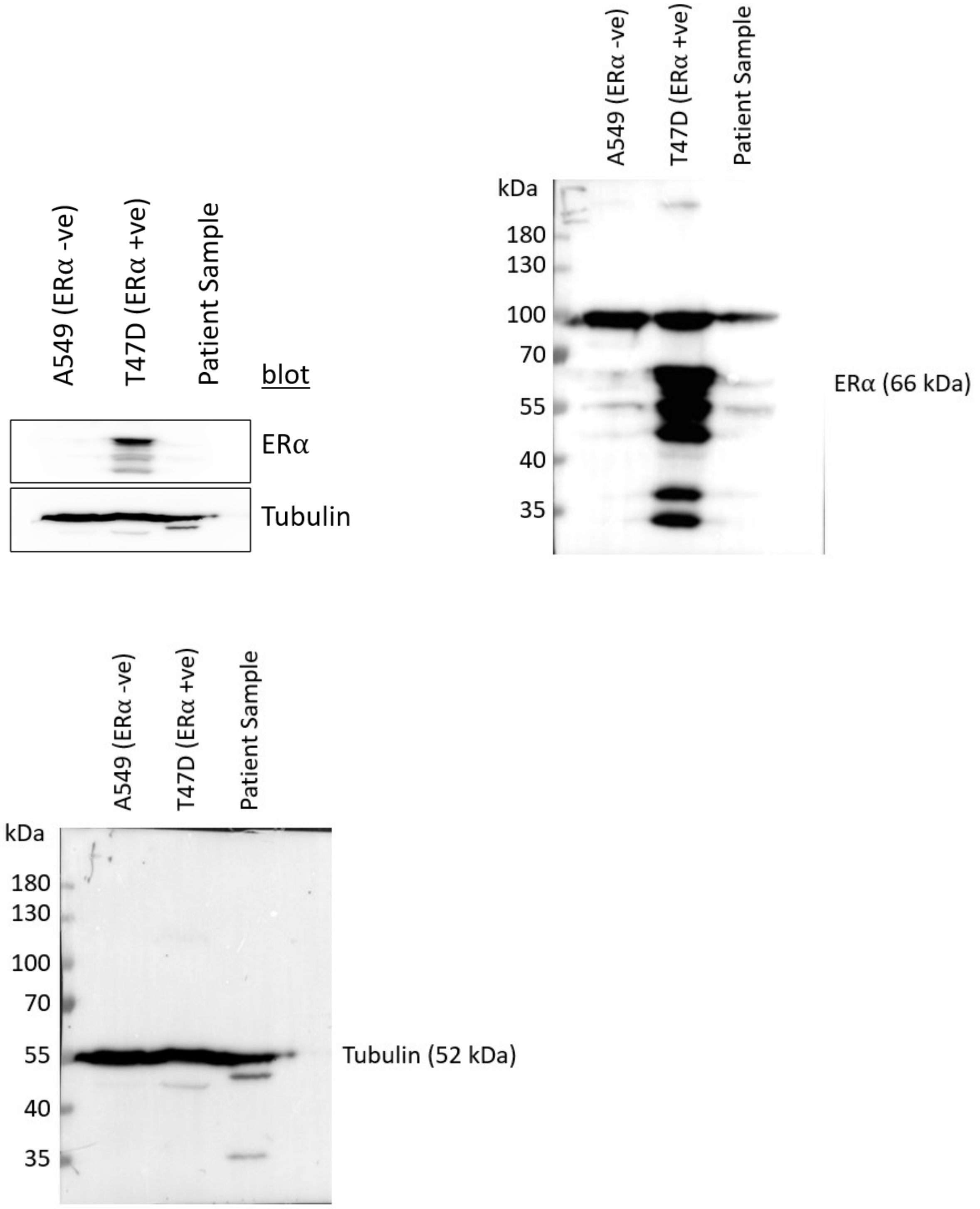


**Supplementary Figure S1. Estrogen receptor α is undetectable in patient P3.** Two breast cancer cell lines, A549 (ERα -ve) and T47D (ERα +ve), have been used as a negative and positive control correspondingly. Cells from A549, T47D and tumour tissue of patient P3 were lysed in RIPA buffer. Protein concentrations were measured and adjusted, and 60 µg of protein was loaded per lane. ERα and Tubulin expression were visualized using immunoblotting.

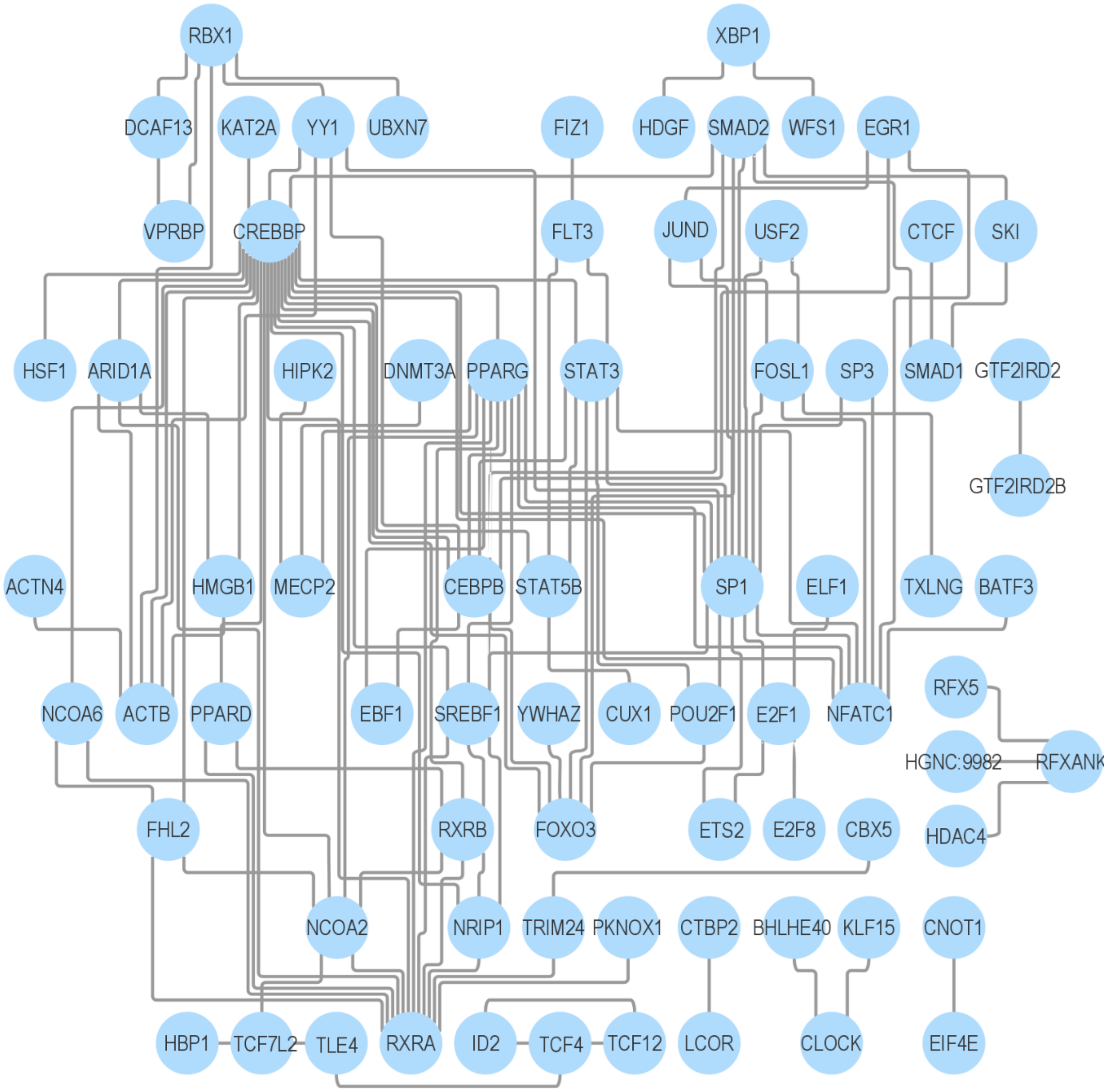


**Supplementary Figure S2**. Network of DNA-binding proteins encoded by genes marked by nucleosome gain at their promoters in tumour tissues. The network was constructed with Cytoscape 3.9.1 (Shannon et al. 2003) using STRING database (Szklarczyk et al. 2023), selecting entries with gene ontology term “DNA-binding”.

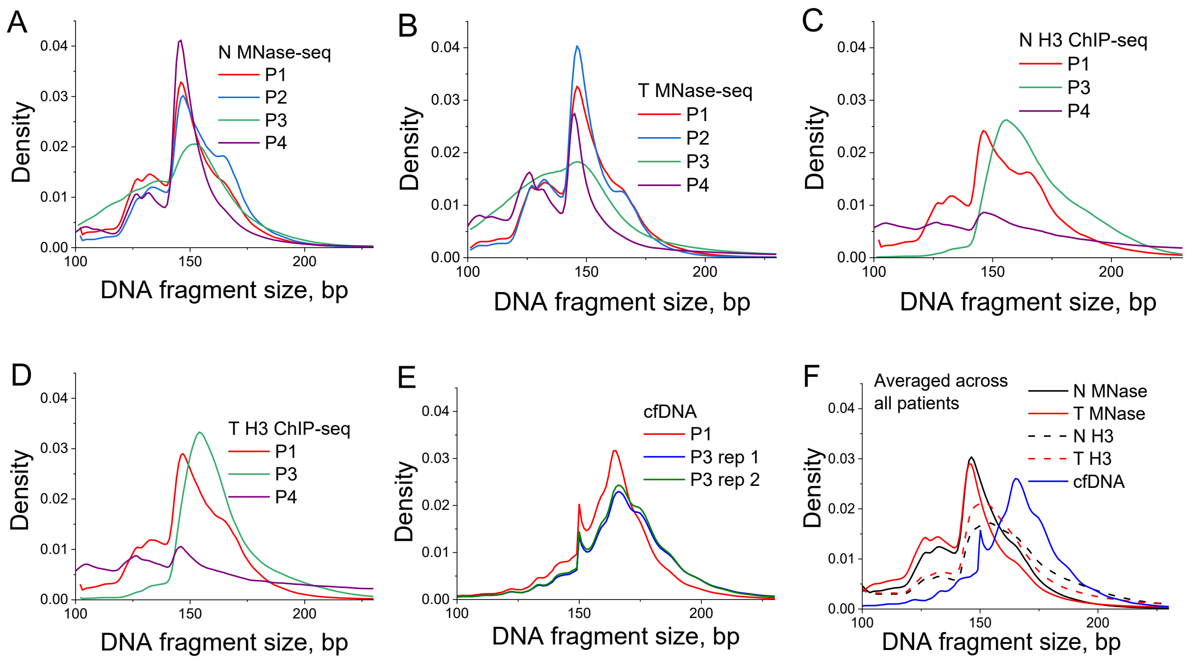


**Supplementary Figure S3. Distribution of DNA fragments protected from digestion in individual patients and averaged across all patients.** A) and B) MNase-seq; C) and D) MNase-assisted H3 ChIP-seq; E) cfDNA from blood plasma. N represent normal breast tissue samples, T represents tumour breast tissue. P1, P2, P3, P4 denote different patients. F) Same as A-E, averaged across all patients in each condition.

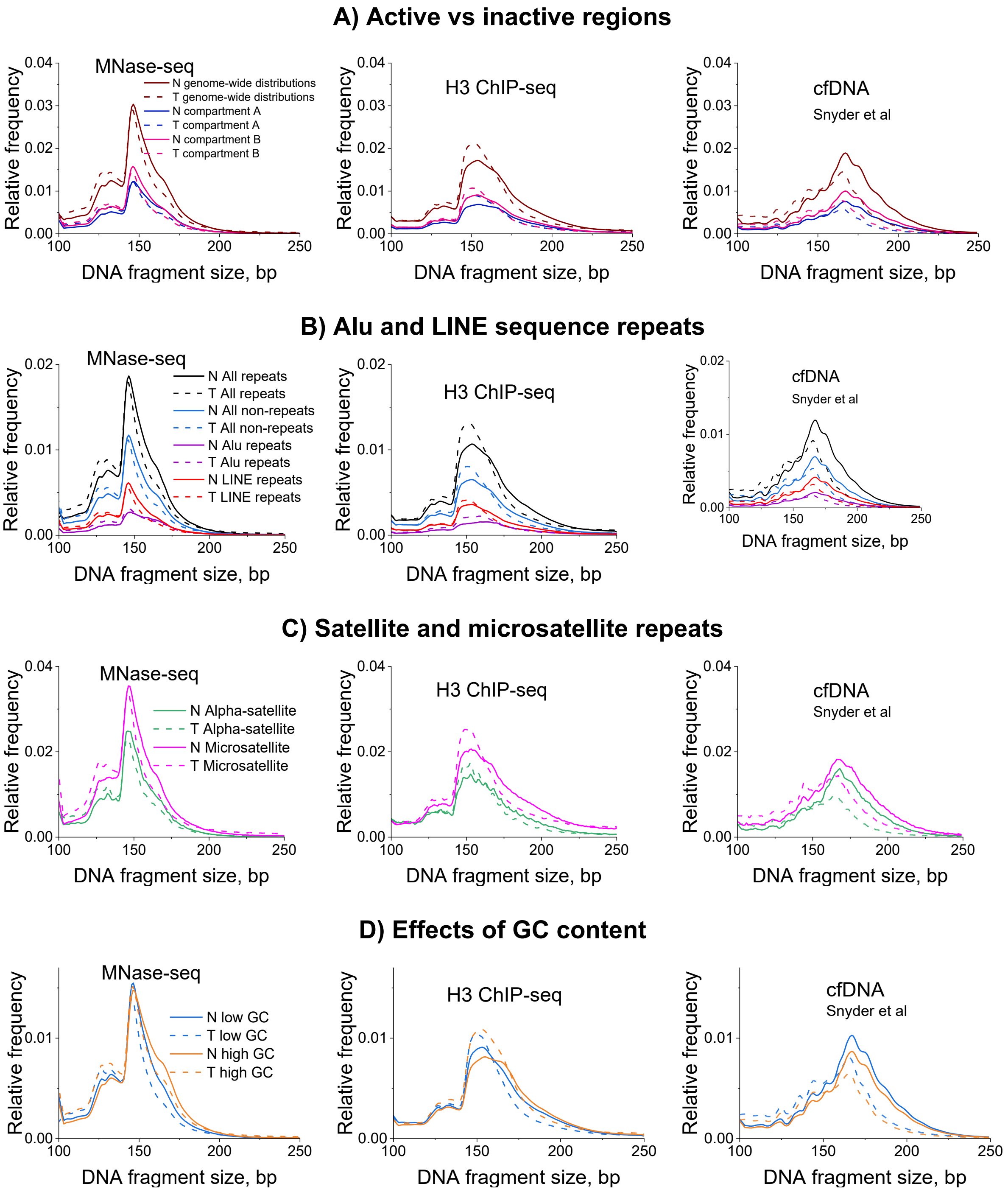


**Supplementary Figure S4. Distribution of DNA fragments protected from digestion, averaged across all samples in a given condition based on MNase-seq, MNase-assisted H3 ChIP-seq from this study and cfDNA sequencing from Snyder et al (Snyder et al. 2016).** A) DNA fragment size distributions inside A and B compartments in MCF7 cells and genome-wide. B-C) DNA fragment size distributions inside different types of repetitive and non-repetitive genomic regions indicated in the figure. D) DNA fragment size distributions inside 10-kb genomic regions with high and low GC content. The solid lines represent normal samples and dashed line represents tumour samples.

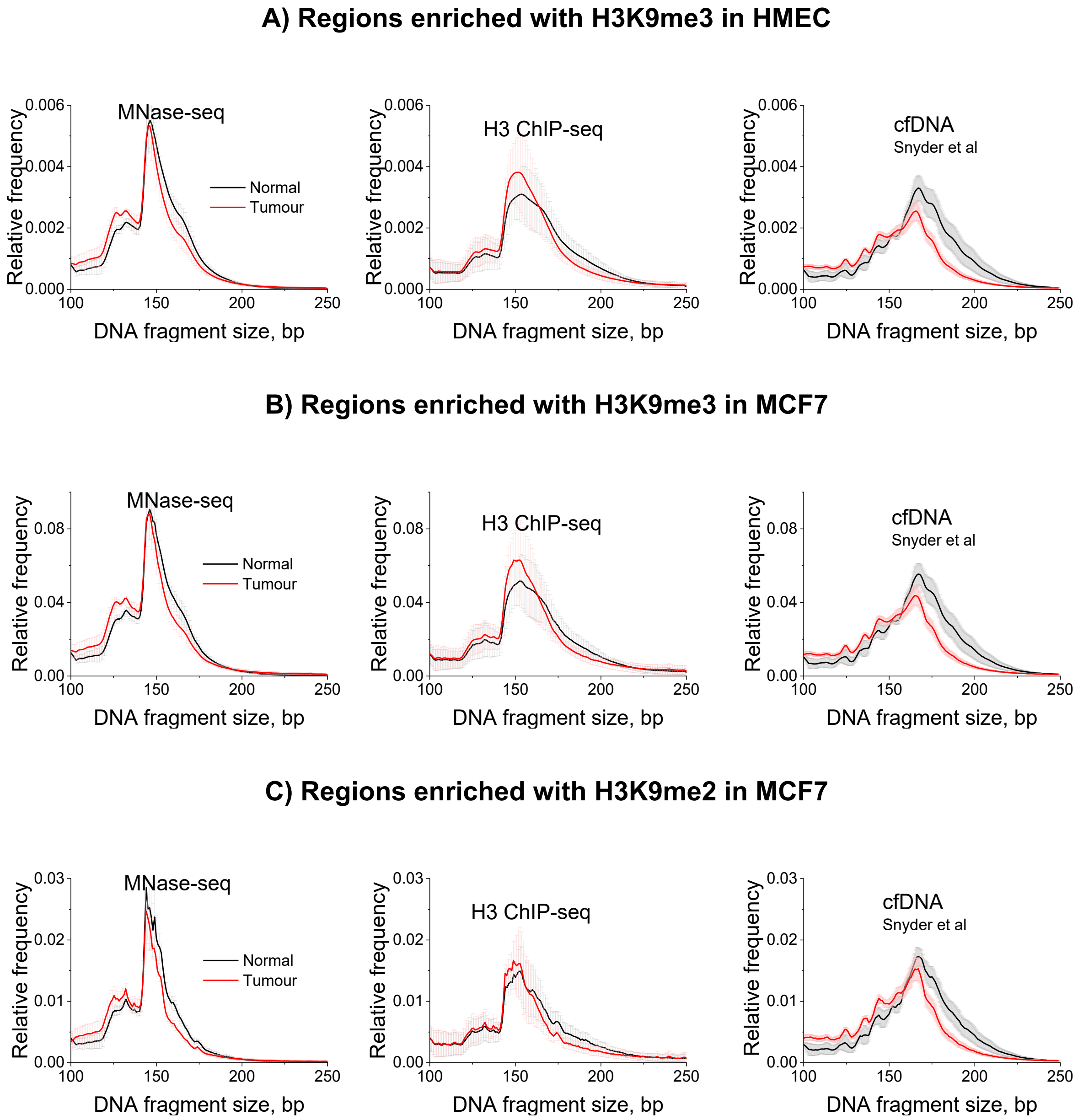


**Supplementary Figure S5. Distribution of DNA fragment sizes protected from digestion inside genomic regions enriched with H3K9me2/3 histone modifications.**

A) DNA fragment size distribution based on MNase-seq, MNase-assisted H3 ChIP-seq and cfDNA sequencing (Snyder et al. 2016) inside regions enriched with H3K9me3 in HMEC cells (healthy breast cells) (Zhang et al. 2020).

B) DNA fragment sizes based on MNase-seq, MNase-assisted H3 ChIP-seq and cfDNA sequencing inside regions enriched with H3K9me3 in MCF-7 cells (Dunham et al. 2012).

C) DNA fragment sizes based on MNase-seq, MNase-assisted H3 ChIP-seq and cfDNA sequencing inside regions enriched with H3K9me2 in MCF-7 cells (Dunham et al. 2012). The solid line corresponds to the averaged profile across all samples in a given condition. The red/grey clouds around the average line show the corresponding standard deviation of averaging.

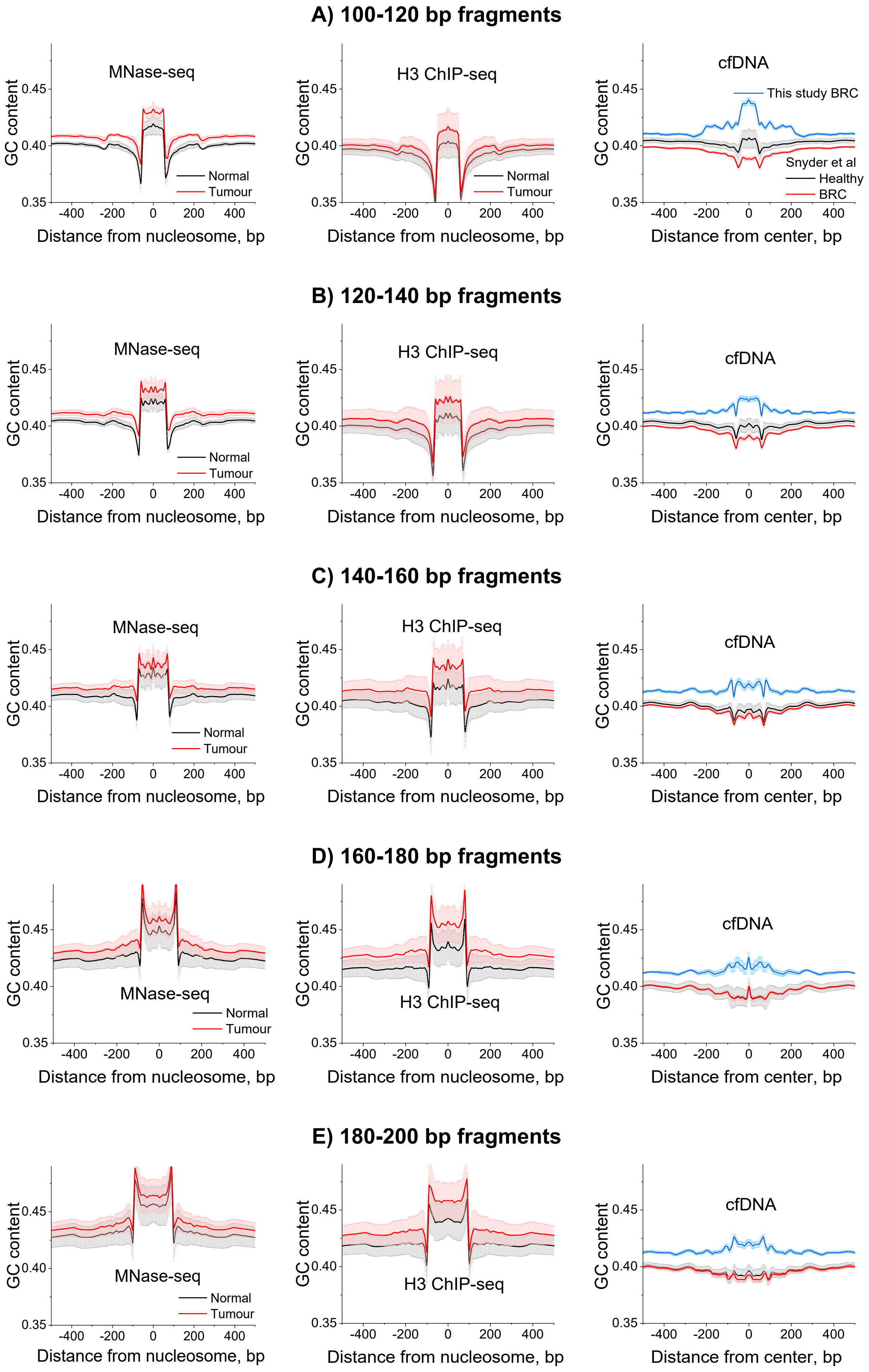
A) 100-120 bp fragments
MNase-seq
H3 ChIP-seq
cfDNA
This study BRC
Snyder et al
Healthy
BRC
Normal
Tumour
GC content
0.35
0.40
0.45
-400
-200
0
200
400
Distance from nucleosome, bp
Distance from center, bp
B) 120-140 bp fragments
C) 140-160 bp fragments
D) 160-180 bp fragments
E) 180-200 bp fragments

**Supplementary Figure S6. GC profiles around DNA fragments protected from digestion in normal/breast cancer tissues and cfDNA genome-wide.** Left column: MNase-seq experiments. Middle column: MNase-assisted H3 ChIP-seq. Right column: cfDNA reported by Snyder et al., 2016 (black, healthy controls and red, BRC patients) and cfDNA from this study (BRC patients, blue). DNA fragment sizes were selected in the following groups: 100-120 bp (A), 120-140 bp (B), 140-160 bp (C), 160-180 bp (D), 180-200 bp (E). The solid line corresponds to the averaged profile across all samples in a given condition. The red/grey/blue clouds around the average line show the corresponding standard deviation of averaging.

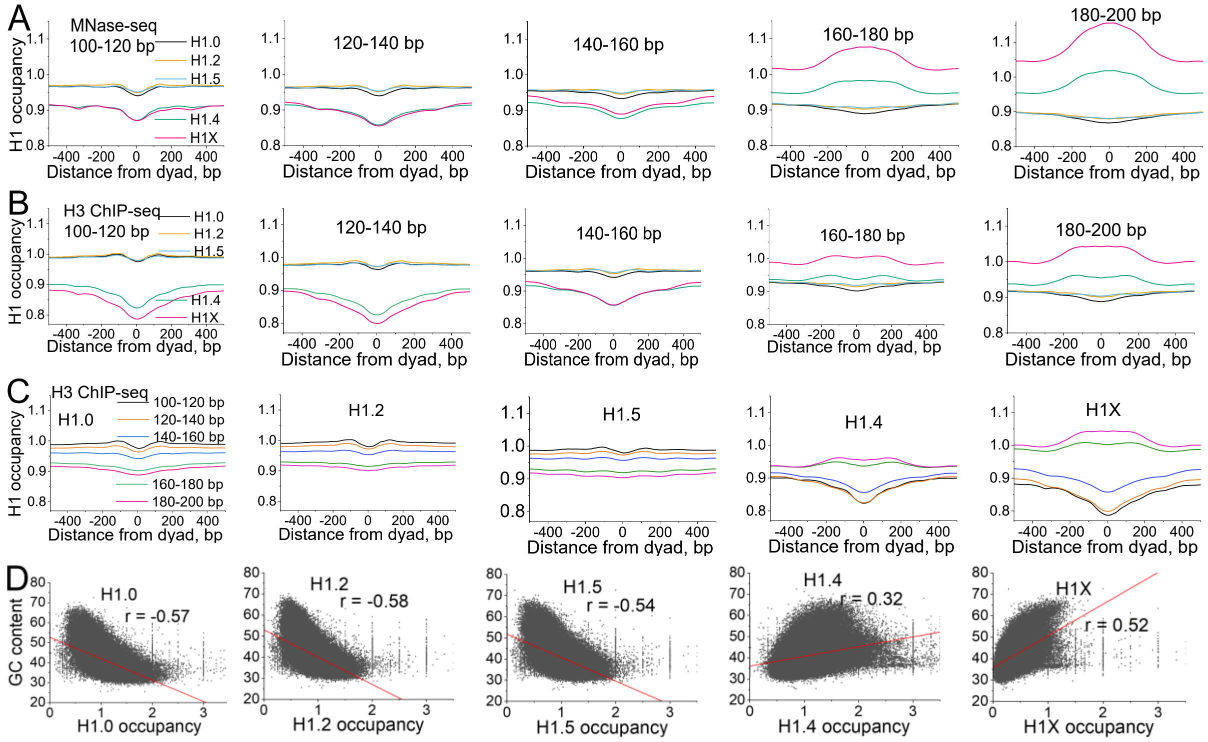


**Supplementary Figure S7**. **Occupancy of linker histone H1 variants around DNA fragments protected from MNase digestion in breast cancer tissues inside genes.**

A) MNase-seq. B) MNase-assisted H3 ChIP-seq. The profiles are shown separately for DNA fragment sizes in the following groups: 100-120 bp, 120-140 bp, 140-160 bp, 160-180 bp,180-200 bp, for the following H1 variants were selected: H1.0 (black), H1.2 (orange), H1.5 (blue), H1.4 (green) and H1X (pink).

C) Aggregate profiles of occupancy of H1 variants around nucleosome dyads calculated for different DNA fragment sizes in MNase-assisted H3 ChIP-seq. The profiles are shown separately for DNA fragment sizes in the following groups: 100-120 bp (black), 120-140 bp (orange),140-160 bp (blue),160-180bp (green) and 180-200 bp (pink).

D) Correlation of occupancy of H1 variants with GC content as in Figure 3A, but calculated inside gene bodies, with each point corresponding to one gene.

**Supplementary Table S1. Clinical information of patients P1 and P2.**

| Patient ID | Age | Meno stats | Tumour size | Lymph nodes | ER | PR | Stage | Grade | Diagnosis |
|---|---|---|---|---|---|---|---|---|---|
| P1 | 84 | Post | 25 | Neg | 7 | 3 | II | 1 | Invasive ductal carcinoma |
| P2 | 60 | Post | 9 | Pos | 8 | 7 | II | 2 | Mixed Invasive Ductal and Lobular Carcinoma |

**Supplementary Table S2.** Numbers of DNA fragments used in the analysis. Sample names are composed of the patient codes, P1, P2, P3, P4; N – normal breast tissue; T – tumour breast tissue; cfDNA – cell-free DNA from the corresponding patient.

| Sample name | Total reads | Uniquely mapped | Reads 120-180 bp |
|---|---|---|---|
| P1 N MNase-seq | 738685119 | 581546291 | 434837001 |
| P1 T MNase-seq | 690415722 | 546197304 | 406009852 |
| P1 N MNase-H3-seq | 799317821 | 593479462 | 405213397 |
| P1 T MNase-H3-seq | 726535624 | 574236019 | 417837382 |
| P1 cfDNA | 605965018 | 305961716 | 455380323 |
| P2 N MNase-seq | 740166473 | 589186960 | 442219740 |
| P2 T MNase-seq | 721399958 | 582290379 | 450655870 |
| P3 N MNase-seq | 719309252 | 552228048 | 367379663 |
| P3 T MNase-seq | 734554908 | 500718271 | 311730314 |
| P3 N MNase-H3-seq | 704821706 | 531125999 | 313518972 |
| P3 T MNase-H3-seq | 825637262 | 578205688 | 386531315 |
| P3 cfDNA 1 | 802041150 | 420816769 | 446372537 |
| P3 cfDNA 2 | 854862937 | 453457506 | 515977495 |
| P4 N MNase-seq | 198922080 | 147359246 | 114522142 |
| P4 T MNase-seq | 176299238 | 107963750 | 65775814 |
| P4 N MNase-H3-seq | 191369339 | 136183830 | 51150348 |
| P4 T MNase-H3-seq | 171247792 | 122174984 | 50702774 |

**Supplementary Table S3. Resequenced reads from the same samples as in Table S2.**

| Sample name | Uniquely mapped | Reads 120-180 bp |
|---|---|---|
| P1 N MNase-seq | 921526763 | 720919673 |
| P1 T MNase-seq | 785058607 | 614106072 |
| P1 N H3 ChIP-seq | 770087909 | 565896992 |
| P1 cfDNA | 590122263 | 455380323 |
| P2 N MNase-seq | 851602991 | 691252262 |
| P2 T MNase-seq | 906470171 | 735504524 |
| P3 N MNase-seq | 729198144 | 506009505 |
| P3 T MNase-seq | 1166043773 | 710577513 |
| P3 cfDNA 1 | 740668098 | 446372537 |
| P3 cfDNA 2 | 808026801 | 515977495 |

**Supplementary Table S4.** Numbers of “stable”, “common”, “shifted”, “lost” and “gained” DNA fragments protected from MNase digestion with sizes 120-180 bp and 160-180 bp.

| | 120-180 bp | 160-180 bp |
|---|---|---|
| **Stable (tumour tissue)** | 418,131,153 | 72,266,802 |
| **Stable (normal tissue)** | 403,536,177 | 107,470,477 |
| **Common** | 308,997,468 | 36,994,858 |
| **Shifted** | 9,460,589 | 6,529,484 |
| **Lost** | 57,210 | 2,310,979 |
| **Gained** | 49,964 | 337,940 |

**Supplementary Table S5. Genome-wide NRL values in normal and tumour breast tissues in the samples from this study.**

| Sample ID | NRL in normal tissue, bp | Standard error, bp (normal) | NRL in tumour tissue, bp | Standard error, bp (tumour) |
|---|---|---|---|---|
| P1 MNase | 191.7 | 0.3 | 192.0 | 0.5 |
| P2 MNase | 200.2 | 0.6 | 193.6 | 0.7 |
| P3 MNase | 203.6 | 0.3 | 197.0 | 0.5 |
| P4 MNase | 199.0 | 0.8 | 192.0 | 0.8 |
| P1 H3 | 198.7 | 0.3 | 191.9 | 0.4 |
| P3 H3 | 204.2 | 0.6 | 195.9 | 0.5 |
| P4 H3 | 199.8 | 2.2 | 191.8 | 2.2 |

**Supplementary Table S6. Genome-wide NRL values in cfDNA samples from this study**

| Sample ID | NRL, bp | Standard error, bp |
|---|---|---|
| P1 cfDNA | 193.9 | 0.5 |
| P3 cfDNA replicate 1 | 192.1 | 0.8 |
| P3 cfDNA replicate 2 | 192.3 | 0.3 |

**Supplementary Table S7. Genome-wide NRL values in human embryonic stem cells (hESCs) calculated based on the MNase-seq dataset GSE49140 by Yazdi et al (Yazdi et al. 2015).** The corresponding accession numbers in the Short Read Archive (SRA) are indicated in the table.

| SRA sample ID | NRL, bp | Standard error, bp |
|---|---|---|
| SRR942485 | 194.8 | 1.6 |
| SRR942489 | 195.2 | 1.0 |
| SRR942490 | 195.6 | 0.6 |
| SRR942492 | 195.1 | 0.8 |
| SRR942493 | 194.6 | 0.6 |
| SRR942497 | 194.8 | 0.9 |
| SRR942498 | 193.4 | 1.3 |

**Supplementary Table S8. Genome-wide NRL values in MCF-7 cells calculated based on MNase-seq datasets GSE77526* and GSE51097 by Shimbo et al (Shimbo et al. 2013).**

| Sample ID | NRL, bp | Standard error, bp |
|---|---|---|
| GSE51097, replicate 1 | 192.1 | 0.2 |
| GSE51097, replicate 2 | 194.9 | 0.2 |
| GSE77526, 0.1 U MNase | 192.6 | 1.3 |
| GSE77526, 0.2 U MNase | 192.0 | 1.2 |
| GSE77526, 0.4 U MNase | 194.9 | 1.9 |
| GSE77526, 0.8 U MNase | 194.6 | 0.6 |
| GSE77526, 1.6 U MNase | 194.8 | 0.4 |
| GSE77526, 4 U MNase | 195.4 | 0.7 |
| GSE77526, 8 U MNase | 194.8 | 0.4 |
| GSE77526, 16 U MNase | 195.8 | 1.0 |

*This dataset contains samples digested with different amounts of MNase units indicated in the table

**Supplementary Table S9. Genome-wide NRL values in T47D cells calculated based on the MNase-seq dataset GSE74308 by Lavender et al (Lavender et al. 2016).** The corresponding accession numbers in the Short Read Archive (SRA) are indicated in the table.

| SRA sample ID | NRL, bp | Standard error, bp |
|---|---|---|
| SRR2774668 | 187.65 | 0.2 |
| SRR2774669 | 189.03 | 1.2 |
| SRR2774670 | 187.09 | 1.2 |
| SRR2774671 | 187.92 | 1.2 |
| SRR2774672 | 189.03 | 1.2 |

### Supplementary References


Dunham I Kundaje A Aldred SF Collins PJ Davis CA Doyle F Epstein CB Frietze S Harrow J Kaul R et al. 2012. An integrated encyclopedia of DNA elements in the human genome. *Nature* **489**: 57-74.

Lavender CA, Cannady KR, Hoffman JA, Trotter KW, Gilchrist DA, Bennett BD, Burkholder AB, Burd CJ, Fargo DC, Archer TK. 2016. Downstream Antisense Transcription Predicts Genomic Features That Define the Specific Chromatin Environment at Mammalian Promoters. *PLoS genetics* **12**: e1006224.

Shannon P, Markiel A, Ozier O, Baliga NS, Wang JT, Ramage D, Amin N, Schwikowski B, Ideker T. 2003. Cytoscape: a software environment for integrated models of biomolecular interaction networks. *Genome Res* **13**: 2498-2504.

Shimbo T, Du Y, Grimm SA, Dhasarathy A, Mav D, Shah RR, Shi H, Wade PA. 2013. MBD3 localizes at promoters, gene bodies and enhancers of active genes. *PLoS genetics* **9**: e1004028.

Snyder MW, Kircher M, Hill AJ, Daza RM, Shendure J. 2016. Cell-free DNA comprises an in vivo nucleosome footprint that informs its tissues-of-origin. *Cell* **164**: 57-68.

Szklarczyk D, Kirsch R, Koutrouli M, Nastou K, Mehryary F, Hachilif R, Gable AL, Fang T, Doncheva NT, Pyysalo S et al. 2023. The STRING database in 2023: protein-protein association networks and functional enrichment analyses for any sequenced genome of interest. *Nucleic Acids Res* **51**: D638-D646.

Yazdi PG, Pedersen BA, Taylor JF, Khattab OS, Chen YH, Chen Y, Jacobsen SE, Wang PH. 2015. Nucleosome Organization in Human Embryonic Stem Cells. *PLoS One* **10**: e0136314.

Zhang J, Lee D, Dhiman V, Jiang P, Xu J, McGillivray P, Yang H, Liu J, Meyerson W, Clarke D. 2020. An integrative ENCODE resource for cancer genomics. *Nature communications* **11**: 1-11.
